# *MYC* controls metastatic heterogeneity in pancreatic cancer

**DOI:** 10.1101/2021.01.30.428641

**Authors:** Ravikanth Maddipati, Robert J. Norgard, Timour Baslan, Komal S. Rathi, Amy Zhang, Pichai Raman, Jason R. Pitarresi, Maximilian D. Wengyn, Taiji Yamazoe, Jinyang Li, David Balli, Michael J. LaRiviere, Ian W. Folkert, Ian D. Millstein, Jonathan Bermeo, Erica L. Carpenter, Scott Lowe, Christine Iacobuzio-Donahue, Faiyaz Notta, Ben Z. Stanger

## Abstract

The degree of metastatic disease varies widely amongst cancer patients and impacts clinical outcomes. However, the biological and functional differences that drive the extent of metastasis are poorly understood. We analyzed primary tumors and paired metastases using a multi-fluorescent lineage-labeled mouse model of pancreatic ductal adenocarcinoma (PDAC) – a tumor type where most patients present with metastases. Genomic and transcriptomic analysis revealed an association between metastatic burden and gene amplification or transcriptional upregulation of *MYC* and its downstream targets. Functional experiments showed that MYC promotes metastasis by recruiting tumor associated macrophages (TAMs), leading to greater bloodstream intravasation. Consistent with these findings, metastatic progression in human PDAC was associated with activation of MYC signaling pathways and enrichment for MYC amplifications specifically in metastatic patients. Collectively, these results implicate MYC activity as a major determinant of metastatic burden in advanced PDAC.

## Main Text

Tumor heterogeneity, most commonly studied in a primary disease setting, is a critical driver of phenotypic diversity, culminating in metastatic, lethal cancers (*1–5*). In most cancers, prognosis and therapeutic decisions are defined by the presence or absence of metastasis. However, tumor heterogeneity is increasingly being questioned at the level of metastatic disease, with recent studies in several cancer types suggesting that metastasis is not a binary phenotype but rather a disease spectrum ranging from oligo- (limited) to polymetastatic (widespread) disesase (*6–8*). Heterogeneity in the manifestation of metastatic disease can guide decisions on use of local regional vs. systemic therapies with emerging evidence of its importance in clinical outcome (*9–11*). Despite its clinical significance, the mechanisms that underlie this spectrum of metastatic states remains unclear and largely understudied.

Pancreatic ductal adenocarcinoma (PDAC) represents a disease entity well suited for the study of metastasis, as most PDACs present with metastatic disease that is associated with dismal prognosis (*12*). Genomic studies have comprehensively catalogued core mutations responsible for primary tumor development in PDAC (e.g. *KRAS*, *TRP53*, *CDKN2A*, and *SMAD4*), paving the path for genomic investigations of metastatic disease for the identification of metastasis promoting alterations. Indeed, recent sequencing studies as well as functional analysis in model systems have associated genomic amplification in muntant KRAS allele with progression from non-metastatic (stage III) to metastatic disease (stage IV) state (*13*). However, genetic factors mediating metastasic heterogeneity in patients and, importantly, the downstream cellular mechanisms, remain largely undefined (*14–18*). Furthermore, it is unclear whether metastasis-associated alterations can influence the tumor microenvironment (TME) – and immune cells in particular – which have been shown to strongly influence metastatic behavior (*19–31*). Therefore understanding the interplay between genetic alterations that influence metastatic behavior and the tumor biology that promotes it – in a cell autonomous and/or non-cell autonomous manner – is crucial for understanding metastasis as a distrinct disease state and critical for the development of more effective treatments.

One barrier to understanding metastatic heterogeneity has been a paucity of model systems that capture this natural variation and allow for direct assessments of paired primary tumors and metastases *in vivo*. This has limited our ability to define the factors intrinsic to primary tumors that influence the extent of metastatic spread. We previously developed an autochthonous model of PDAC – the KPCX model – that employs multiplexed fluorescence-based labeling to track the simultaneous development of multiple primary tumor cell lineages and follow them as they metastasize (*32*). Importantly, this technique facilitates confirmation of lineage relationships *in vivo*, such that primary tumor clones with substantial metastatic potential can be distinguished from those having poor metastatic potential. Here, we show that this system recapitulates the variation in metastatic burden found in human PDAC and use it to dissect molecular and cellular features contributing to metastatic heterogeneity.

## Results

### Metastatic burden is variable in human and murine PDAC

While the vast majority of PDAC patients have metastases (principally liver and lung), the number of metastases is highly variable from patient to patient (*33, 34*). Importantly, data regarding metastases have largely been obtained at autopsy and thus confounded by varying treatment histories and reseeding due to end-stage disease (*4*). Thus, we first sought to characterize the burden of metastases in treatment-naïve patients. To this end, we performed a retrospective analysis of initial CT scans from 55 patients newly diagnosed with metastatic (stage IV) PDAC at the University of Pennsylvania (Fig. 1a). The total number of lesions in the lung and liver were counted by examining both coronal and sagittal planes for both organs and binned into groups of ten, revealing a wide distribution of metastatic burden (Fig. 1b). K-means clustering identified two metastatic subgroups: a Met^low^ subgroup (≤10 metastases, 25/55) and a Met^high^ subgroup (>10 metastases, 30/55) (Fig. 1b, Fig. S1a). Primary tumor size, age, sex, and race were not correlated with differences in metastatic burden (Fig. 1c, Fig. S1b). However, having a greater number of metastases was associated with worse overall survival (Fig. 1d). Thus, even among patients with stage IV PDAC, metastatic burden is variable and correlates with clinical outcome.

**Figure 1:**
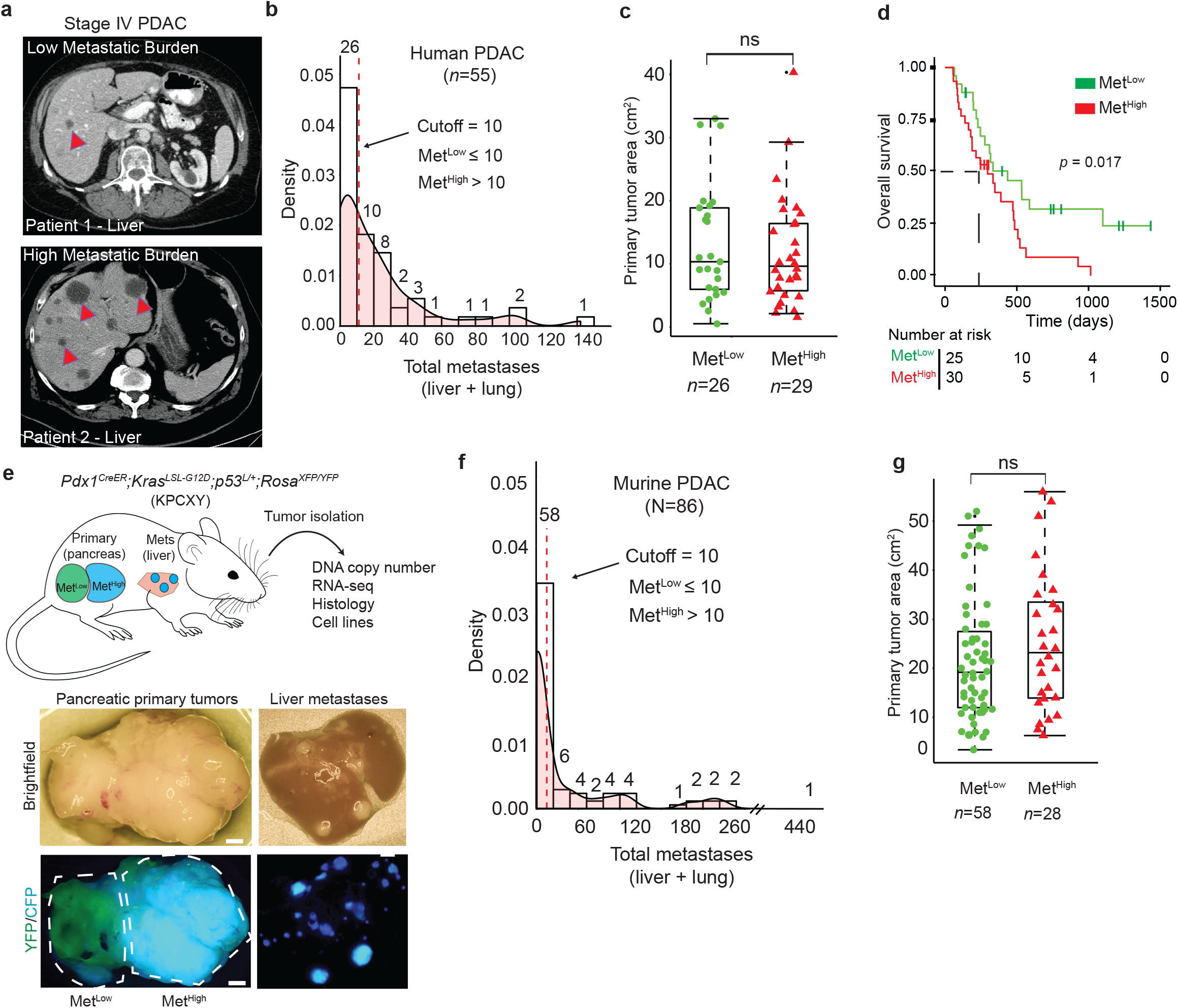
Advanced pancreatic tumors exhibit intertumoral differences in their propensity for metastasis. **a.** CT imaging of human PDAC liver metastasis demonstrating heterogeneity in metastatic burden in Stage IV disease. Arrowheads indicate solitary metastasis in the top panel and selected metastases in the bottom panel **b.** Density plot and histogram showing the distribution of total (liver and lung) metastases enumerated from CT scans of human Stage IV PDAC at the time of diagnosis (n=55). Values above each histogram bar represent the number of patients in each group. The vertical dotted line (red) represents the cutoff between Met^Low^ tumors (≤ 10 mets) and Met^High^ tumors (>10 mets) determined by k-means clustering. **c.** Quantification of tumor area (based on tumor dimensions from largest cross-sectional plane on imaging) comparing Met^Low^ and Met^High^ cases from the cohort in (**b**). **d.** Overall survival analysis of the cohort in (**b**). **e.** Top: Schematic view of the KPCXY model, showing multiple primary tumors distinguishable by color arising in the pancreas with matched metastases in the liver. Bottom: Representative fluorescent stereomicroscopic images showing a YFP+ tumor adjoining a CFP+ tumor in the pancreas (left) and liver metastases derived from the CFP+ tumor in the same animal (right). **f.** Density plot and histogram showing the distribution of total (liver and lung) metastases enumerated at autopsy of KPCXY mice. Values above each histogram bar represent the number of tumors giving rise to the indicated number of metastases, based on color (n=85 tumors from 30 KPCXY mice). The vertical dotted line (red) represents the cutoff between Met^Low^ tumors (≤ 10 mets, N=58) and Met^High^ tumors (>10 mets, N=28) determined by k-means clustering. **g.** Quantification of tumor area comparing Met^Low^ and Met^High^ tumors from the cohort in (**f**). Statistical analysis by Student’s unpaired t-test with p-values indicated (ns, not significant). Box-whisker plots in (**c**) and (**f**) indicate mean and interquartile range. Scale bar (**e**) = 1mm.

We hypothesized that the differences in metastatic burden seen in human PDAC may also be present in autochthonous murine models. To test this, we used the KPCXY model – in which Cre-mediated recombination triggers expression of mutant **K**ras^G12D^ and **p**53^R172H^ in the pancreatic epithelium along with **Y**FP and confetti (**X**) lineage tracers (Fig. 1e, Methods) – to measure metastatic heterogeneity in a cohort of tumor bearing mice. By exploiting the multi-color features of the KPCX model, we previously showed that these mice harbor (on average) 2-5 independent primary tumor clones; importantly, the clonal marking of tumors with different fluorophores makes it possible to infer the lineages of primary tumors with different metastatic potential (*32*). In our earlier work with this model, we noted that in most tumor-bearing animals – even those with multiple primary tumors – metastasis to the liver and lung were driven by a single tumor clone (Fig. 1e, Fig. S1c). This suggested that tumor cell-intrinsic factors strongly influence the metastatic behavior of a tumor, even within a single animal.

To quantify differences in metastatic burden, we examined a panel of mice with at least 2 uniquely labeled fluorescent tumors where most metastases could be attributed to a specific tumor on the basis of color (Fig. 1e, Fig. S1c). A total of 85 primary tumors from 30 mice were examined, and gross metastases to the liver and lung arising from each tumor were then quantified by stereomicroscopy (Methods). Murine PDACs exhibited a wide distribution of metastatic burden, with a pattern resembling that of the human disease (Fig. 1f). Similarly, k-means clustering grouped murine samples into a low metastasis subgroup (≤10 metastases, 58/85) and a high metastasis subgroup (>10 metastases, 27/85), which we similarly refer to as the Met^low^ and Met^High^ subgroups, respectively (Fig. 1f, Fig. S1d). As with the human disease, neither primary tumor size nor tumor cell proliferation correlated with metastatic burden (Fig. 1g, Fig. S1e). Thus, the KPCXY model recapitulates the intertumoral metastatic heterogeneity seen in human PDAC and provides a unique experimental model for comparing highly metastatic and poorly metastatic tumor clones.

### Individual tumor lineages in KPCXY mice correspond to clones with distinct somatic copy number profiles

Although primary KPCXY tumors were easily distinguishable based on the expression of a distinct fluorophore, each tumor could have arisen via the clonal expansion of a single cell or through fusion of multiple tumors which happened to share the same color. Somatic copy number alterations (SCNAs) have been shown to provide an unambiguous picture of genomic heterogeneity and lineage relationships between primary tumors and matching metastases in human disease (*35*). Consequently, we performed copy number analysis via genome sequencing on a set of 20 primary tumors, including multi-regional sampling on a subset of the tumors where sufficient tissue was available (9 tumors with 2-4 regions sampled per tumor) (Fig. 2a, Table S1). Tumors bearing different colors exhibited unique DNA copy number profiles, indicating that they arose independently (Fig. 2b, Fig. S2a)(*36*). By contrast, multi-regional sampling of monochromatic tumors revealed shared copy number alterations, indicating that all subregions within a given tumor (defined by color) shared a common ancestral lineage (Fig. 2c, Fig. S2b). In addition, subregion-specific alterations were also observed, suggesting that subclonal heterogeneity is also present in each tumor (Fig. 2c, Fig. S2b). These results suggest that the monochromatic tumors observed in KPCXY mice are clonal in origin and continue to undergo subclonal evolution during tumor progression.

**Figure 2:**
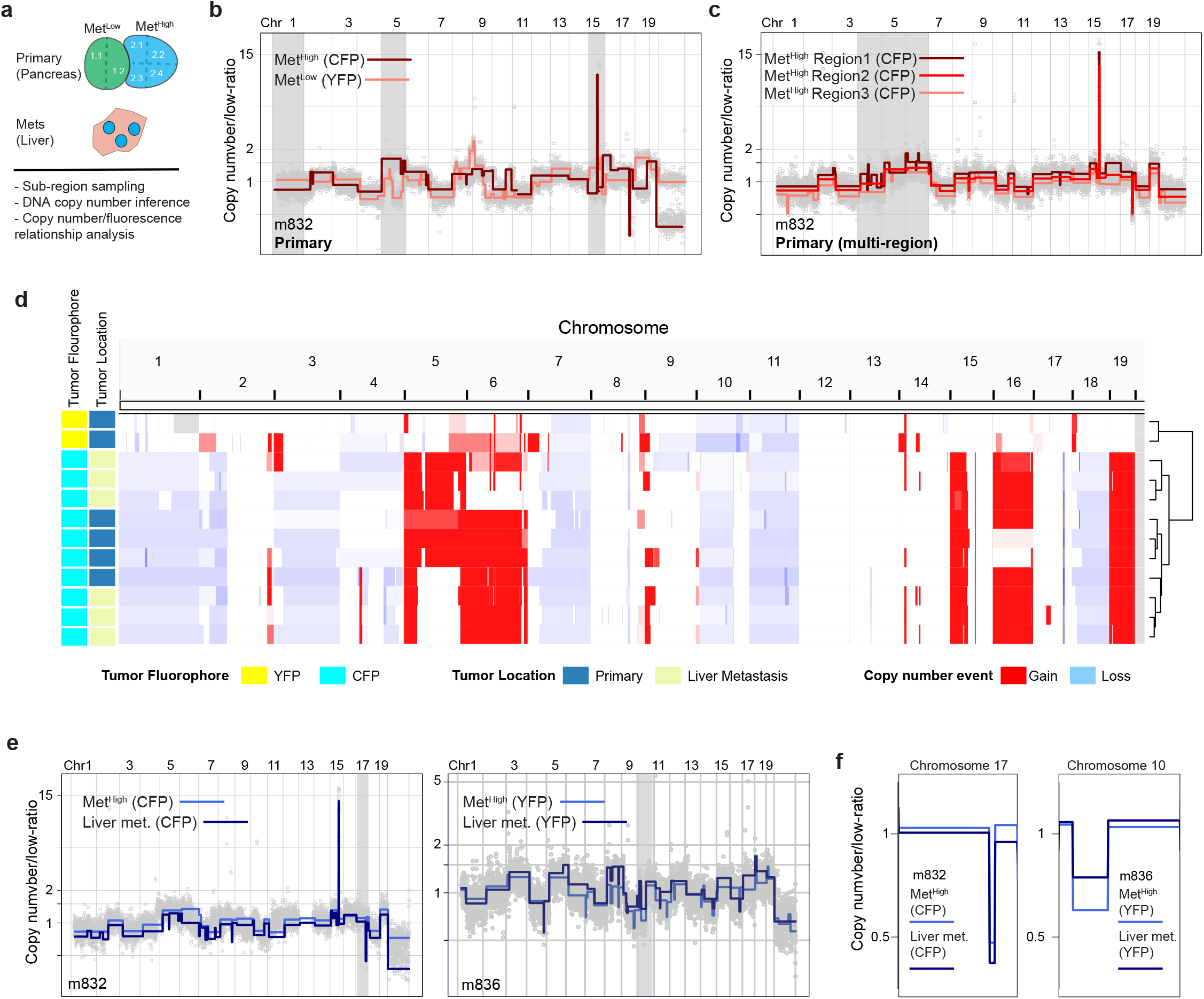
Somatic copy number alteration analysis confirms fluorescence based lineage relationships and reveals genetic heterogeneity in in paired primary pancreatic tumors and liver metastasis. **a.** Schematic representation of KPCXY pancreatic tumor and matching liver metastasis with multi-region sampling for copy number sequence analysis. **b.** Representative genome-wide copy number profiles of Met^High^ (CFP+ fluorescence) and Met^Low^ (YFP+ fluorescence) tumors from mouse 832 (m832) as depicted in Fig 1e. Gray shading denotes alterations that are unique to the Met^High^ (CFP+) tumor. Y-axis illustrate normalized read count values (low-ratio) which are directly proportional to genome copy number at a given chromosomal location. The copy number profiles are centered around a mean of 1 with gains and deletions called for segments with values higher and lower than the mean, respectively (Methods). **c.** Representative genome-wide copy number profiles of three sub-sampled tissue regions of the Met^High^ (CFP+) primary tumor from m832. Gray shading denotes alterations that are found heterogeneously from multi-region sequencing of the primary tumor. **d.** Genome-wide heatmap with hierarchal clustering based on copy number alterations of matched primary and metastatic samples profiled from m832. **e.** Representative genome-wide copy number profiles of fluorescently matched primary and metastatic tissue from two profiled mice (m832-left panel and m836-right panel) illustrating the shared clonal genetic lineage. **f.** Zoom-in chromosomal views of copy number alterations with distinguishing breakpoint patterns supporting shared genetic lineage. Panels are ordered as in (**e**).

To ascertain the lineage relationships between primary tumors and metastases, we compared DNA copy number profiles between liver metastases and primary tumors within a given mouse. This revealed that primary tumors and metastases of the same color shared common DNA copy number profiles across the dataset, confirming on a genetic basis the fluorescence based lineage relationships (Fig. 2d-f, Fig. S2c). As most lung metastasis were microscopic and difficult to isolate by dissection, they were not included in the molecular analysis. Together, these results indicate that the lineage history of metastases can be inferred by color and genomic analysis, allowing primary tumors with high vs. low metastatic potential to be unambiguously classified.

### Genomic and transcriptional analyses identify *Myc* as a potential driver of metastatic phenotypes

We next sought to examine the molecular differences that distinguish primary tumors with high vs. low metastatic potential. We began by examining large scale (mega-base level as well as chromosome wide) SCNAs in 20 Met^High^ and Met^Low^ primary tumor samples. This analysis revealed largely similar genome-wide copy number patterns between Met^high^ and Met^low^ primary tumors with key PDAC associated genes, such as loss-of-heterozygosity (LOH) at *Cdkn2a/b* and *Trp53* as well as chromosomal gain of *Kras* occurring at similar frequencies (Fig. S3a). Thus, KPCXY tumors exhibit frequent copy number alterations in canonical PDAC genes, but these alterations do not account for the variation in metastatic behavior between Met^high^ and Met^low^ tumors.

We next asked whether other factors (genomic and/or transcriptional) may be acting to enhance metastasis in the Met^High^ group. Focal amplifications in driver oncogenes – *Cdk6* and *Yap* in breast cancer and mutant *Kras* in PDAC – have been linked to the acquisition of metastatic competence(*13, 14, 37, 38*). Consistent with prior studies, we observed focal amplicons at genomic regions encoding *Cdk6*, *Yap,* and *Kras* in our tumors (Fig. 3a)(*13, 14, 37–39*). However, in contrast to these amplifications, which occurred at equal frequencies in Met^High^ and Met^Low^ tumors, focal high amplitude amplifications in *Myc* were found in 42.8% (3/7) of Met^High^ tumors compared to 7.6% (1/13) of Met^Low^ tumors (Fig. 3b). Thus, *Myc* amplifications are enriched in Met^high^ tumors. In all cases, these amplifications were maintained in paired metastases (Fig. S3b). In addition, RNA-seq analysis demonstrated significantly higher levels of *Myc* transcripts in Met^High^ tumors and metastases compared to Met^Low^ tumors (Fig. 3c); overall, *Myc* was the third-most significantly upregulated gene in Met^High^ tumors compared to Met^Low^ tumors (Fig. 3d). Gene set enrichment analysis (GSEA) of the differentially expressed genes between Met^High^ and Met^Low^ tumors identified MYC and E2F signatures as highly enriched, along with other signatures that have been implicated in PDAC metastasis including unfolded protein response, oxidative phosphorylation, and hypoxia (Fig. 3e)(*31, 40, 41*). Moreover, MetaCore transcription factor enrichment analysis identified MYC as the TF most significantly associated with genes overexpressed in Met^High^ tumors (Fig. 3f), and Ingenuity Pathway Analysis placed *Myc* at the center of the interactome generated by these differentially expressed genes (Fig. S4). Collectively, these results demonstrate a strong association between a tumor’s metastatic behavior and the abundance and/or activity of *Myc* at the genomic and transcriptional levels.

**Figure 3:**
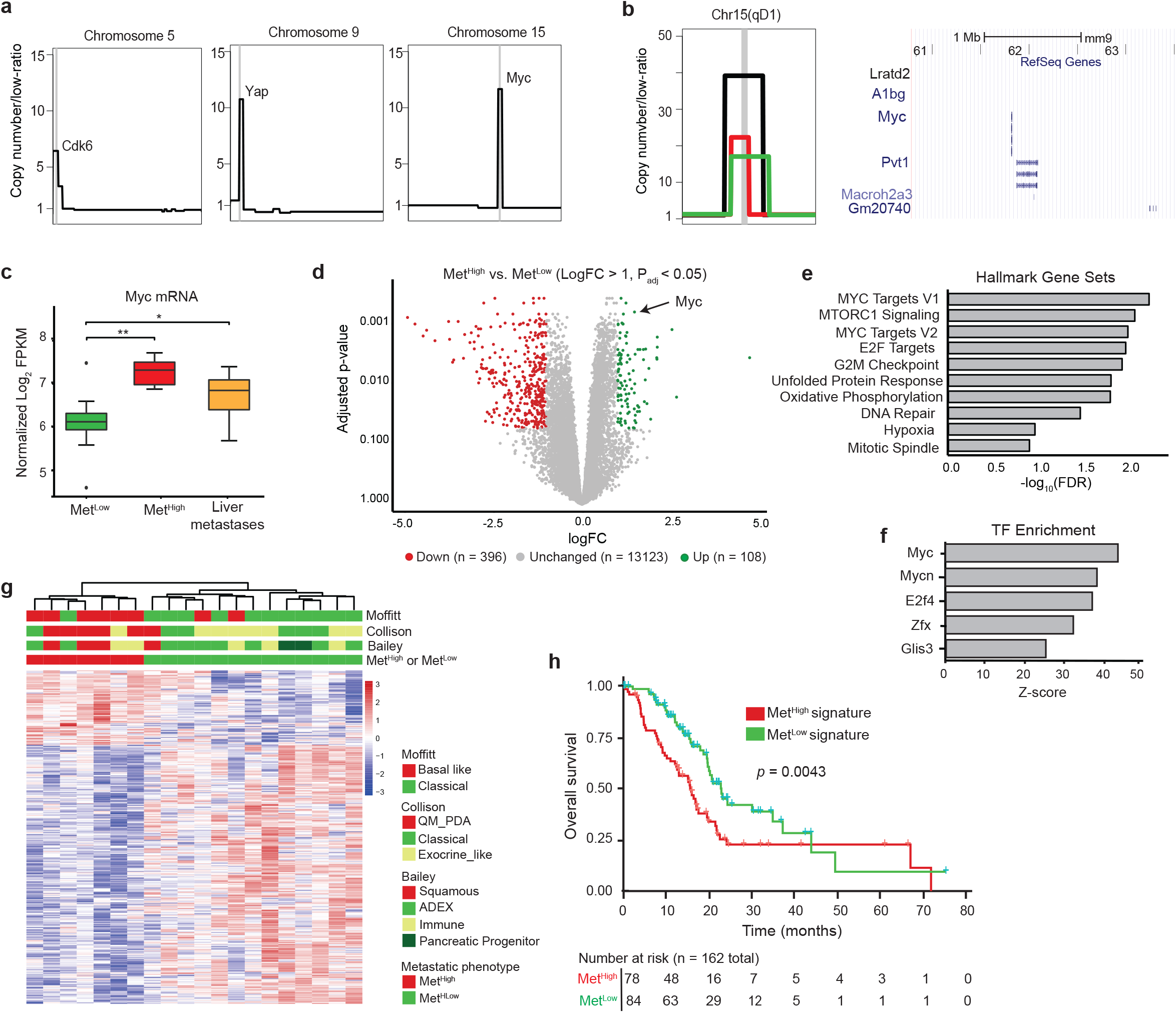
The Met^high^ phenotype is associated with focal, high amplitude *Myc* amplifications and elevated expression. **a.** Schematic representation of focal amplifications identified in profiled primary tumors. Vertical gray line denotes the location of amplicon and likely driver gene **b.** (Left) Zoomed in schematic representation of three identified *Myc* amplicons in Met^High^ tumors illustrating the focal and high-amplitude nature of the event. Each event (amplicon) is illustrated by a different colored segment line. The shared amplified region between the different amplicons is denoted by the chromosomal cytoband top of panel and illustrated in UCSC genome browser view (Right) with RefSeq Genes, including *Myc*, illustrated. **c.** Box-and-whisker plot showing *Myc* mRNA levels in Met^High^ tumors (n=7) and paired metastases (n=34) compared to Met^Low^ tumors (n=13). **d.** Volcano plot illustrating genes meeting cutoffs for differential expression (logfold-change >1, Padj <0.05) between Met^High^ and Met^Low^ tumors (n=20 tumors used in the comparison). Genes upregulated in Met^High^ tumors are highlighted in green, and genes upregulated in Met^Low^ tumors are highlighted in red. **e.** Top ten Hallmark Gene Sets identified as enriched in Met^High^ tumors compared to Met^Low^ tumors using all DEGs (adj. p < 0.05). **f.** Top five transcription factor binding sites enriched in DEG in Met^High^ tumors compared to Met^Low^ tumors (adj. p< 0.05) identified by Metacore prediction software. **g.** Heatmap showing unsupervised clustering of differentially expressed genes (logfold-change >1, Padj <0.05) between Met^High^ and Met^Low^ tumors (n=20) and their association with PDAC transcriptional subtypes previously reported by Collison(*42*), Moffitt(*15*), and Bailey(*16*). **h.** Kaplan-Meier analysis showing overall survival of PDAC patients in the TCGA cohort stratified into those with a Met^High^ signature (red line) versus those with a Met^Low^ signature (blue line). Signature based on DEGs with absolute logfold-change > 0.58 and Padj <0.05 (736 up and 1036 down regulated genes) Statistical analysis in (**c**) was performed by Wilcoxon test (*, p=3.9×10^−4^; **, p=5.3×10^−5^). Box and whiskers represent median mRNA expression and interquartile range. Statistical analysis in (**h**) was performed by log-rank test.

Human PDAC can be grouped into two main transcriptomic subtypes – a well-differentiated classical/exocrine-like/progenitor (classical) subtype and a poorly differentiated squamous/quasi-mesenchymal/basal (basal-like) subtype (*15, 16, 18, 42*). We found that Met^High^ tumors were strongly associated with basal-like PDACs, in line with their more aggressive behavior (Fig. 3g). Likewise, applying murine Met^Low^ and Met^High^ signatures (see Methods) to human TCGA data predicted a worse survival – indicative of disease recurrence – for patients with a Met^High^ signature (Fig. 3h). These data indicate that murine Met^High^ tumors correspond to the more aggressive subtypes of human PDAC.

### A panel of cell lines that preserve the Met^Low^ and Met^High^ phenotypes

To understand the mechanisms underlying these different metastatic properties, we generated a panel of cell lines from six Met^High^ tumors and five Met^Low^ tumors. Consistent with the parent *in vivo* tumors, *Myc* gene expression and *Myc* protein levels were higher in the Met^High^ lines compared to the Met^Low^ lines (Fig. 4a-b). SCNA analysis in these cells lines found that they retained the majority of the genomic alterations found in the matched primary samples, including *Myc* amplifications (Fig. S5a-b). Furthermore, *Myc* amplifications were not found in any of the cell lines whose tumors were originally characterized as non-*Myc* amplified, indicating that *in vitro* culture does not select for this specific copy number alteration. Importantly, elevations in *Myc* mRNA and protein were observed in both the *Myc* amplified and non-amplified Met^High^ lines, suggesting that elevated *Myc* expression is a stable phenotype of these cells in culture.

**Figure 4:**
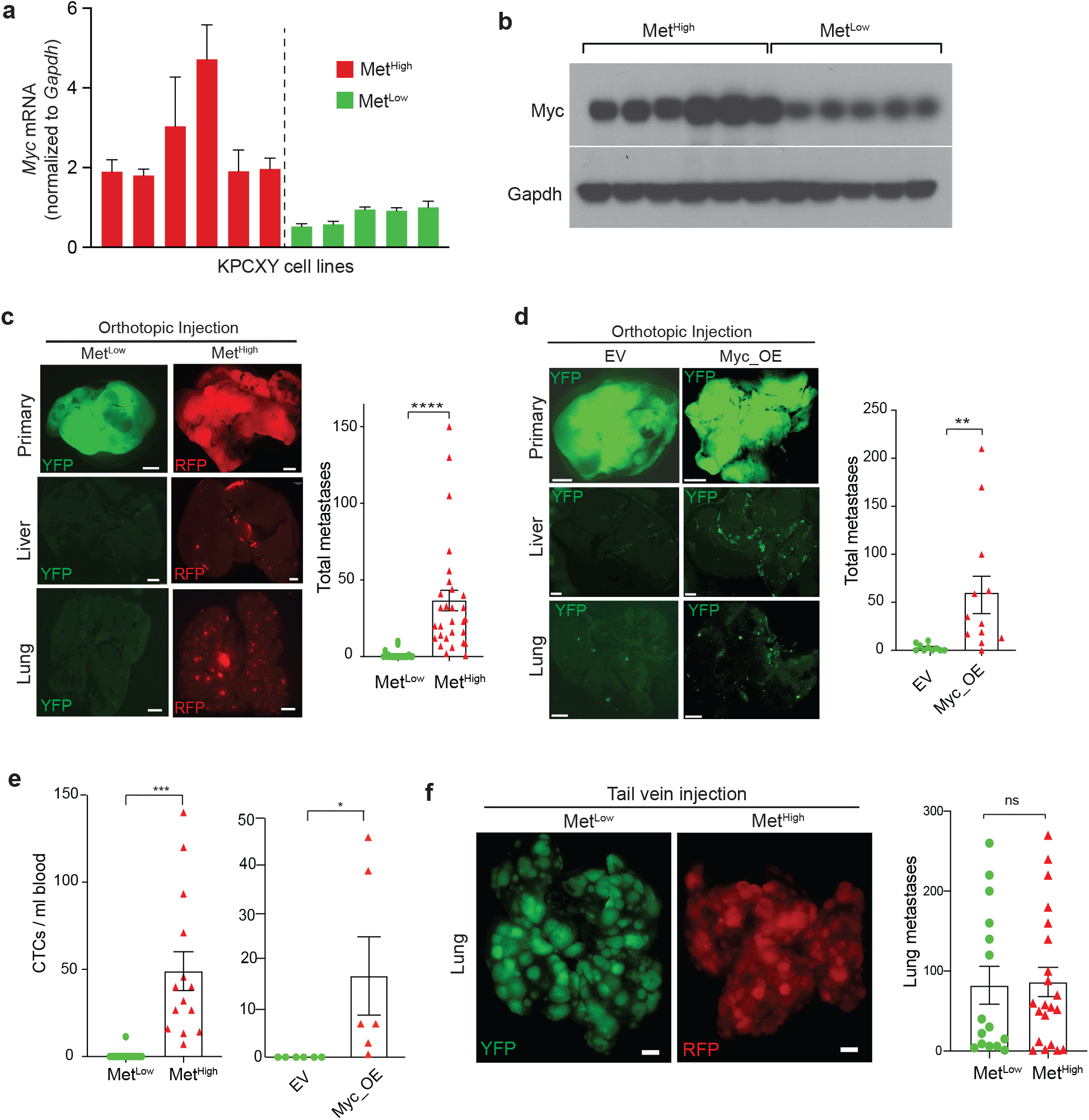
*Myc* regulates metastasis by enhancing tumor cell intravasation. **a.** Bar graph showing *Myc* mRNA levels in cell lines derived from Met^High^ and Met^Low^ tumors, normalized to Gapdh. N= 6 Met^High^ and 5 Met^Low^ cell lines **b.** Western blot showing corresponding MYC protein levels in cell lines derived from Met^High^ and Met^Low^ tumors described in (**a**). **c.** Representative fluorescent images of primary tumors and associated liver and lung metastasis following orthotopic transplantation of the cell lines in (**a**) and (**b**) into NOD.SCID mice. The bar graph shows the total number of metastases (liver and lung) counted following orthotopic transplantation of 5 Met^Low^ cell lines or 5 Met^High^ cell lines (pooled data from N=49 mice in total). **d.** Representative fluorescent image of primary tumors, liver, and lung metastases following orthotopic transplantation of Met^Low^ cell lines that were stably transduced with either a *Myc* overexpression construct (Myc-OE) or empty vector (EV). The bar graph shows the total number of metastases (liver and lung) counted following orthotopic transplantations of Myc-OE or EV cells. Data were pooled from 4 independent Met^Low^ lines transduced with either the MYC-OE or EV construct transplanted into 12 NOD.SCID (for the Myc-OE cells) or 10 NOD.SCID mice (for the EV cells). **e.** Quantification of circulating tumor cells (CTCs) in arterial blood derived from the orthotopic tumors depicted in (**c**) (n=27 mice examined) and d (n= 12 mice examined). **f.** Representative fluorescent images of lung metastasis following tail vein injection of cell lines derived from the Met^Low^ and Met^High^ primary tumor clones. The bar graph shows the total number of lung metastases counted following tail vein injection of 5 Met^Low^ cell lines or 5 Met^High^ cell lines (pooled data from N= 36 mice in total). Statistical analysis by Student’s unpaired t-test with significance indicated (*, p=0.0152; **, p = 0.013; ***, p= 0.0008; **** p < 0.0001; ns, not significant). Error bars indicate SEM (**c-f**). Scale bar = 1 mm (**c-d,f**).

To investigate the metastatic properties of the Met^High^ and Met^Low^ lines *in vivo*, we performed orthotopic implantation of 5 Met^High^ and 5 Met^Low^ lines into the pancreas of NOD.SCID mice and examined distant organs for evidence of metastasis. Although the weights of Met^High^ and Met^Low^ tumors were not significantly different (Fig. S6a), Met^High^ tumors gave rise to 28-fold more liver and lung metastases compared to Met^Low^ tumors (Fig. 4c). Consistent with the cell line expression differences, the orthotopic Met^High^ tumors expressed higher levels of *Myc* compared to Met^Low^ tumors (Fig. S6b). To further confirm that differences in *Myc* expression were sufficient to drive the metastatic phenotype, we introduced a *Myc* overexpression construct into 4 Met^Low^ lines and generated orthotopic tumors (Fig. S6c). *Myc* overexpression led to a dramatic (22-fold) increase in liver and lung metastases (Fig. 4d) which could not be accounted for by the modest increase in tumor weight (Fig. S6d). Thus, cell lines derived from spontaneously-generated Met^High^ and Met^Low^ tumors retain their metastatic phenotypes upon implantation.

### MYC promotes tumor cell intravasation through the recruitment of tumor associated macrophages

To form distant metastases, cancer cells must navigate a series of events collectively referred to as the “metastatic cascade.” These events include (i) intravasation into the bloodstream or lymphatics, (ii) survival in the circulation, (iii) extravasation from the vessel, and (iv) growth and survival at the distant site(*43*). To determine the step(s) at which *Myc* was exerting its prometastatic effects, we began by measuring the number of circulating tumor cells (CTCs) in orthotopically-implanted Met^High^ and Met^Low^ tumors and in Met^Low^ tumors engineered to overexpress *Myc* (Myc-OE). Remarkably, CTCs arising from Met^High^ and Myc-OE tumors were 38-fold and 17-fold more abundant than those arising from Met^Low^ tumors (Fig. 4e), far greater than the approximately 2-fold increase in tumor weight resulting from Myc overexpression (Fig. S6d). Next, we performed a tail vein metastasis assay, which bypasses the invasion step by introducing tumor cells directly into the bloodstream and measuring lung metastases. Surprisingly, in contrast to the orthotopic tumor experiment, there was no difference in the number of metastases between Met^High^ and Met^Low^ lines (Fig. 4f). Taken together, these data suggest that Met^High^ tumors achieve a higher metastatic rate principally by promoting cancer cell invasion into the circulation which can be driven by increased *Myc* expression.

Beyond activation of tumor intrinsic programs, *Myc* can also affect tumor phenotypes by altering the tumor immune microenvironment (TiME)(*44–46*). Thus, we sought to determine if differences in *Myc* levels between Met^High^ and Met^Low^ tumors were associated with distinct TiMEs. To this end, we examined the immune composition of parental primary tumors by staining for markers of immune cells previously implicated in metastasis of PDAC and other cancers. While Met^High^ and Met^Low^ tumors had a similar degree of neutrophil infiltration, Met^High^ tumors had lower numbers of CD3+ T-Cells and were highly enriched for F4/80+ macrophages (Fig. 5a). In particular, there was an increased abundance of macrophages positive for Arg1 (Fig. 5b), a marker of alternatively activated tumor associated macrophages (TAMs)(*47*). Thus, compared to Met^Low^ tumors, the TiME of Met^High^ tumors contains an increased number of TAMs.

**Figure 5:**
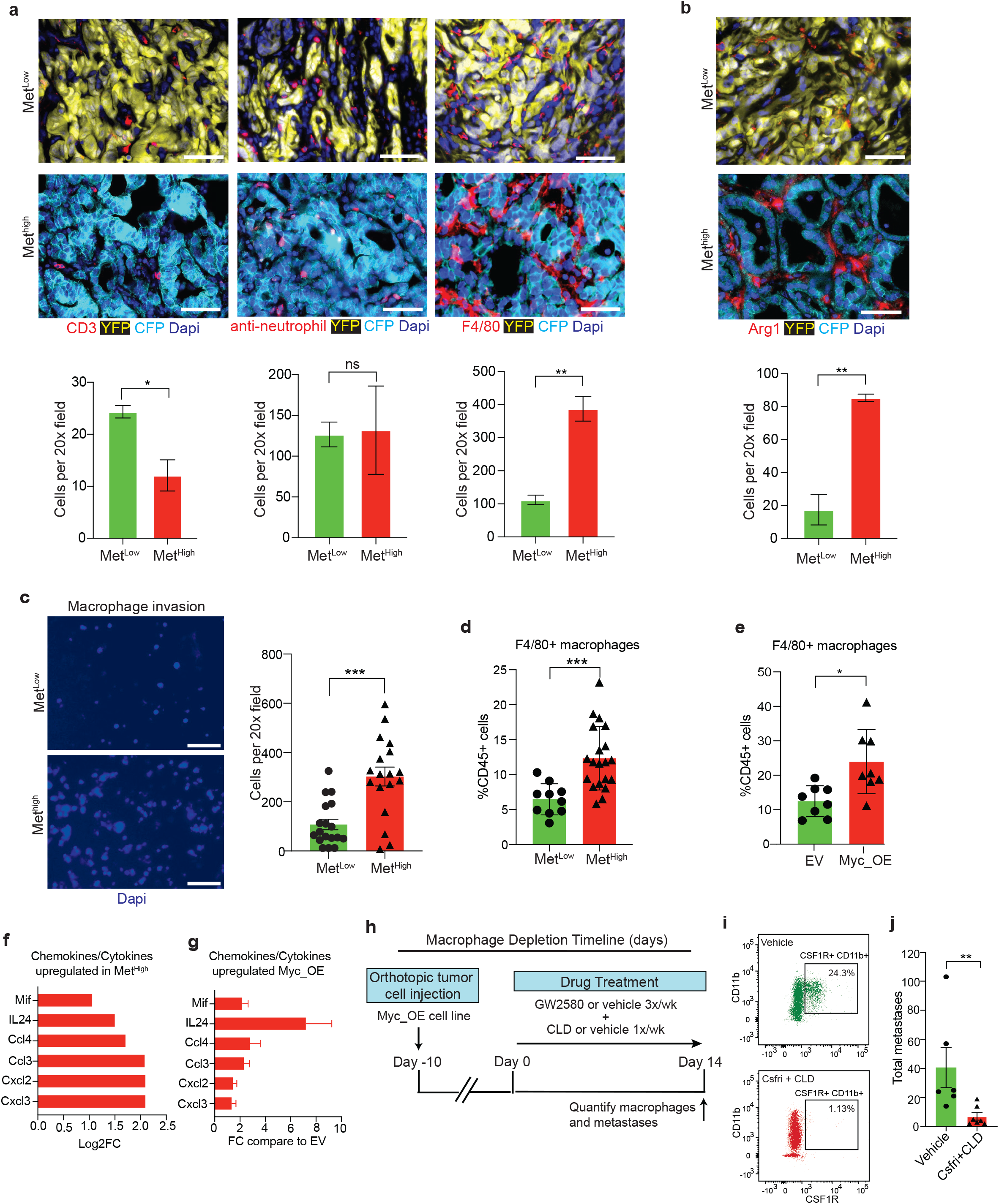
Myc recruits pro-metastatic macrophages to the TME. **a-b.** Representative immunofluorescence images (top) and quantification (bottom) of T-cells (CD3+), Neutrophils (anti-neutrophil antibody+), macrophages (F4/80)+, and Arg1+ macrophages in primary KPCXY tumors categorized as Met^Low^ or Met^High^ (n=3 mice for each subgroup and 4-5 random fields of view analyzed). **c.** Representative immunofluorescence images (left) and quantification (right) of macrophages that have migrated across a transwell filter following co-culture with Met^High^ or Met^Low^ tumor cells (n= 2 Met^Low^ and 2 Met^High^ cell lines used, 3 replicates per cell line with 3 20X images taken per transwell; each dot represents quantification of an independent image). **d.** Quantification of tumor infiltrating macrophages (as a percentage of total CD45+ cells) in Met^Low^ or Met^High^ subcutaneous tumors assessed by flow cytometry (n=5 Met^High^ cell lines and 3 Met^Low^ cell lines; 2 NOD.SCID mice examined per cell line with 2 tumors per mouse; each dot represents an independent tumor). **e.** Quantification of tumor infiltrating macrophages (as a percentage of total CD45+ cells) in Myc-OE or control (EV) subcutaneous tumors assessed by flow cytometry (n=2 Myc-OE cell lines and 2 EV cell lines; 2 NOD.SCID mice examined per cell line with 2 tumors per mouse; each dot represents an independent tumor). **f.** Differentially expressed chemokines/cytokines with elevated expression (logfold-change >1, Padj <0.01) between Met^High^ and Met^Low^ tumors (as described in Fig 3d) **g.** Bar graph showing fold change in chemokines/cytokine mRNA levels listed in (f) comparing Myc-OE to EV cell lines. Data representative of 2 independent cell lines. **h.** Schematic outline of the macrophage depletion experiment. Mice were orthotopically implanted with Myc-OE cell lines (n=2 independent cell lines) and after 10d were treated with a combination of CSFR inhibitor (GW2580) and liposomal clodronate (CLD) or vehicle. Metastases were quantified 14d later. **i.** Representative flow cytometry dot plots showing relative abundance of CSFR1+CD11b+ macrophages in the circulation following treatment with GW2580+CLD or vehicle. **j.** Quantification of total metastases (liver and lung) following the macrophage depletion strategy outlined in (**h**) (n=6 control mice and N=7 GW2580+CLD mice; each dot represents an independent mouse). Statistical analysis by Student’s unpaired t-test with significance indicated (*, p<0.05; **, p<0.005; ***, p<0.0001; ns, not significant). Error bars indicate SEM (**a-e, h**). Scale bars = 10 μm (**a-b**) and 50 μm (**c**).

To examine the ability of Met^High^ and Met^Low^ tumors to recruit macrophages in our system, we co-cultured these cell lines with primary bone marrow derived murine macrophages in a transwell migration assay(*39*). Compared to Met^Low^ lines, Met^High^ co-cultures resulted in higher levels of macrophage migration towards the tumor cells (Fig. 5c). Consistent with these *in vitro* results, orthotopic tumors generated from Met^High^ cell lines exhibited greater TAM infiltration than those generated from Met^Low^ cell lines (Fig. 5d). Furthermore, Met^Low^ lines overexpressing *Myc* (Myc_OE) gave rise to tumors with greater TAM infiltration compared to controls (Fig 5e). These results demonstrate that Met^High^ tumors exhibit enhanced macrophage recruitment and implicate *Myc* expression as a driver of macrophage infiltration.

The recruitment of macrophages to tumors can occur in response to factors secreted by tumor cells. To identify potential mediators of macrophage recruitment by Met^High^ tumors, we mined our RNAseq data to identify secreted factors that are differentially expressed between Met^High^ and Met^Low^ tumors (p value <0.01 and logFC >1). This resulted in identification of six cytokines/chemokines upregulated in Met^High^ tumors (Fig 5f). Importantly, MYC overexpression in the Met^Low^ cell lines resulted in upregulation of these factors as well (Fig 5g). Interestingly, each of these factors has been previously implicated in regulating macrophage recruitment in pancreatic and other cancer types (*48–52*). These data suggest that multiple *Myc*-regulated factors likely act in concert to induce macrophage recruitment and metastsis.

To directly test the role of TAMs in metastasis, we generated orthotopic tumors from Met^Low^+Myc_OE lines. After 10 days, we treated mice with a combination of colony stimulating factor receptor inhibitor (CSFRi) to inhibit macrophage migration and liposomal clodronate (CLD) to ablate tissue resident macrophages (Fig. 5f). This regimen was highly effective at depleting macrophages (Fig. 5g), consistent with prior studies (*29, 53, 54*). Macrophage depletion resulted in a 4-6-fold reduction in metastases (Fig. 5h), suggesting that MYC enhances metastatic spread in part by creating a TAM-rich environment in the primary tumor. While previous studies have implicated macrophages in tumor cell infiltration, our work directly links this process to the genomic and transcriptional activation of Myc, which occurs naturally in our model and is subsequently selected for as a driver of metastasis (*21, 26, 28, 52, 53, 55–60*).

### Metastasis in human PDAC is associated with *MYC* gene amplification and elevated expression

Given the finding that genomic and transcriptional variation in *Myc* was associated with metastatic heterogeneity in murine PDAC, we sought to determine whether *MYC* is associated with similar metastatic phenotypes in human PDAC. Since the majority of PDAC samples in the ICGC and TCGA are derived from resected stage I/II tumors (*18*), these datasets provide limited insight into the determinants of metastatic burden. Consequently, we analyzed data from the COMPASS trial cohort (NCT02750657) which is focused on metastatic PDAC patients and utilizes laser capture microdissection (LCM) to enrich for tumor cells prior to whole genome sequencing or RNA-seq (*14, 61*). By comparing primary tumors and metastases, we found that 11.3% (n=17/133) of metastatic tumors were enriched for *MYC* amplifications compared to 1.61% (n=4/244) of resectable disease (Fig. 6a-b; p=7.6e-5, Fishers test). Likewise, advanced tumors (defined as either locally advanced or metastatic) were significantly enriched for *MYC* amplifications (9.22%; n=19/206) compared to resectable tumors having no evidence of metastasis at diagnosis (1.04%; n=2/192) (Fig. S7a; p=1.33e-4). As predicted, amplification was associated with higher levels of *MYC* mRNA (Fig. S7b). *MYC* amplified tumors did not exhibit greater genomic instability compared to non-amplified tumors (Fig. S7c). Moreover, metastases expressed higher levels of *MYC* than primary tumors at the mRNA level (Fig. 6c; p=0.00312). These results indicate that *MYC* amplification and transcriptional upregulation are strongly associated with PDAC metastases.

**Figure 6:**
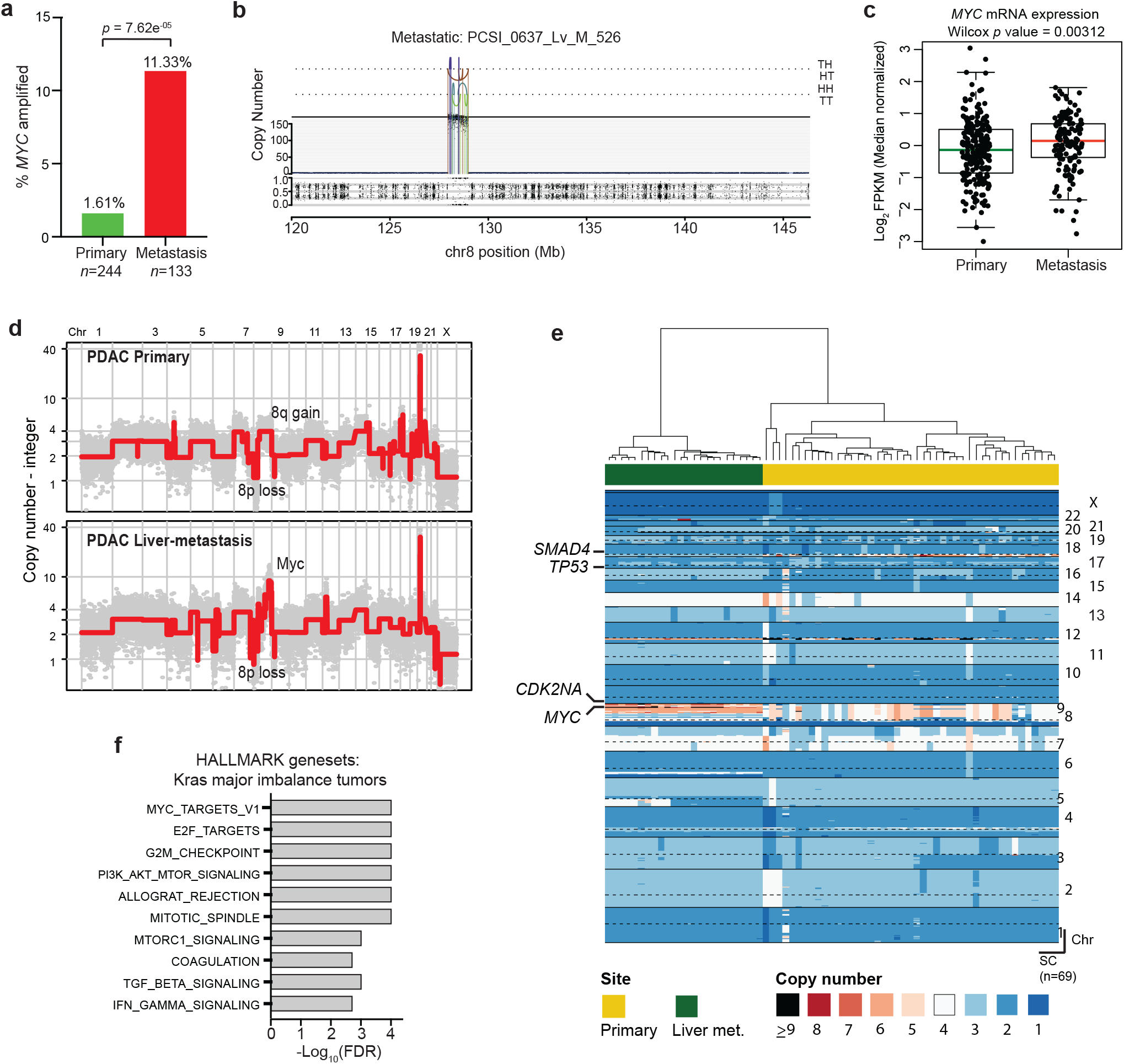
*MYC* amplification and enhanced transcriptional activity are associated with metastasis in human PDAC. **a.** Bar graph showing the relative frequencies of *MYC* amplifications in primary PDAC tumors and metastases from the COMPASS cohort. **b.** Representative plot of chromosome 8 from a metastatic tumor with *MYC* amplification. Orientation of breakpoint junctions from intra-chromosomal rearrangements indicated by TH, HT, HH, and TT where T = tail (3’ end of fragment) and H = head (5’ end of fragment). **c.** Box-and-whisker plot showing *MYC* mRNA levels (FPKM) in primary PDAC tumors and metastases. **d.** Representative genome-wide absolute copy number plots of single cells retrieved from a primary (top panel) and its matched metastasis (lower panel) illustrating acquisition of focal MYC amplification in the metastatic lesion. **e.** Heatmap depiction of cancer single cells sequenced from a matched primary PDAC and its liver metastasis. Color codes indicate absolute copy number in single-cells. Top bar plot depicts tissue site from where single-cells were retrieved. **f.** Gene set enrichment analysis of tumors with a major imbalance of mutant *KRAS* (compared to those with no major imbalance) in the COMPASS cohort. Box-whisker plot in (**c**) indicates mean and interquartile range.

Next, we examined a separate patient cohort (N=20) in which matched primary PDAC tumors and metastases were available for comparison. *MYC* amplifications were common in patients with metastatic disease (35.0%; N=7/20), and these amplifications were retained in the matching metastasis (Fig. S7c), similar to our mouse model. The enrichment of *MYC* amplifications in metastatic samples and the observed retention of the amplification when analyzing matched primary/metastasis samples suggests that amplification and/or transcriptional upregulation of *MYC* in primary PDACs are selected for and retained during tumor metastatic progression. Consistent with this notion, we identified a PDAC patient in whom single cell analysis of a paired primary tumor and metastasis revealed enrichment of a *MYC* amplified subclone in the metastatic lesion compared to the primary (Fig. 6d-e, Fig. S7d). Collectively, these data suggest that enhanced expression and/or genomic amplification of *MYC* is associated with metastatic spread in human PDAC, complementing our findings from the mouse model.

## Discussion

Phenotypic variation, the result of inter- and intra-tumoral heterogeneity arising during tumor progression, has made it challenging to understand the molecular mechanisms underlying tumor spread (*1, 62*). Consequently, the demonstration that certain genes function as “metastasis drivers” – promoting metastasis through mechanisms distinct from their roles in primary tumor growth – has proven elusive (*63*). In this study, we exploited an autochthonous PDAC model with varying degrees of metastatic spread to explore the molecular basis of naturally-arising variation in metastatic burden. This system revealed a strong association between the level of *Myc* – at either the genomic or transcriptional level – and tumor metastasis, a relationship that was also observed in human PDAC samples. *Myc* exerts its pro-metastatic effect at least in part by recruiting pro-invasive TAMs, leading to greater tumor cell intravasation into the bloodstream. These activities are not directly related to *Myc*’s well-described role in primary tumor growth (*46, 64, 65*), as tumors with different levels of *Myc* expression grow at comparable rates despite dramatic differences in metastatic ability.

Prior work by us and others has examined the genetic events associated with PDAC metastasis. In one study, a comparison of matched primary tumors and metastases from four patients failed to reveal nonsynonymous mutations in driver oncogenes that distinguished primary tumors from metastases (*66*). By contrast, examination of DNA copy number changes in mouse and human PDAC revealed a significant association between increased mutant *KRAS* gene dosage and metastatic progression (*13*)(*14*). While our mouse studies did not detect an association between *Kras* focal amplification and metastatic potential, *Kras* gains (via entire gain of chromosome 6) were detected with comparable frequency in both Met^High^ and Met^Low^ tumors, consistent with the ability of both tumor populations to metastasize. Interestingly, the MYC_TARGETS geneset represented the most highly enriched signature in PDAC patients with *KRAS* amplifications or major allelic imbalances (Fig. 6f) mirroring the enrichment of this signature in *MYC* amplified PDAC tumors (Fig. S7f). Signaling through KRAS has long been known to impact MYC expression (*45, 46, 67, 68*), and thus our results are consistent with a model in which elevated MYC activity – as a result of *MYC* and/or *KRAS* amplification, or some other mechanism – enhances metastatic activity. This interpretation is consistent with recent studies of lung, breast, and prostate cancers identifying a link between *MYC* amplification and brain or bone metastasis (*37, 38, 69*).

Although tumor formation in the KPCXY model results from shared founder mutations (*Kras*^*G12D*^ activation and *Trp53* loss) our genomic analysis revealed ongoing somatic events during tumor progression, resulting in heterogeneous patterns of genomic alterations within a given tumor. Such alterations were largely present at the level of copy number gains and losses rather than point mutations or small insertion/deletions. Although the complexity of genomic rearrangements varied between tumors, the degree of genome instability did not correlate with metastatic burden. Thus, the increase in metastasis observed in Met^High^ tumors is not a function of overall SCNA burden but is instead specific to *Myc*.

While subregions within a tumor shared many genomic alterations, consistent with a clonal origin, distinct copy number alterations were also present, suggesting ongoing subclonal evolution. Clonally related metastases exhibited unique (“private”) alterations; however most copy number gains and losses were shared with the parent primary tumor clone, suggesting that they were present prior to dissemination. In this respect, it is noteworthy that *MYC* amplifications in human PDAC were far more common in metastases than primary tumors, including one case in which we were able to trace a metastatic lesion directly to a *MYC*-amplified subclone in the primary tumor. Collectively, these results suggest that subclonal *MYC* amplifications, which have been observed in human primary tumors, provide a selective advantage during metastatic progression (*70, 71*). Given that our analysis identified a *Myc* signature in tumor Met^High^ clones without *Myc* amplifications, and *MYC* amplifications are present in only 11% of human PDAC metastases, other mechanisms for the increased expression of *MYC* mRNA in PDAC metastases are likely to exist.

As one of the best-studied oncogenes, *MYC* has been associated with multiple tumor-promoting activities (*67*). Given MYC’s role in tumor cell growth and proliferation, one possible explanation for our results is that Met^High^ tumors had an earlier onset and/or grew more rapidly, leading to increased metastasis by mass effect. Against this possibility, we found that tumor size and proliferation rates showed no correlation with metastatic burden. Likewise, our Met^High^ and Met^Low^ cell lines exhibited dramatic differences in metastatic ability despite giving rise to primary tumors of comparable size, suggesting that Myc can have a context dependent role wherein it drives tumor growth in early progression and enhances metastatic spread later. Indeed, prior studies have shown that Myc overexpression in the pancreas in the context of tumor initiation does not result in PDAC but rather insulinomas (*72*). By contrast, our work provides strong evidence that Myc overexpression following the establishment of a primary PDAC confers metastatic properties.

Our studies implicate non-cell autonomous mechanisms involving the recruitment of tumor associated macrophages (TAMs) as contributors to metastatic heterogeneity. The ability of TAMs to promote tumor cell invasion is well-documented (*21, 26, 28, 52, 53, 55–60*), a property that is in agreement with our finding that Met^High^ and Myc_OE tumors exhibit enhanced vascular intravasation. Furthermore, MYC expression in tumor cells is known to shape the makeup of the surrounding immune microenvironment, making it more immunosuppressive (*39, 45*). In line with these observations, we find that Met^High^ tumors are enriched for alternatively activated TAMs and have decreased T-cell infiltration – features that favor metastasis (*21, 26, 28, 30, 31, 52, 53, 55–60*). Our data thus support a model wherein stochastically-arising tumor subclones with elevated levels of MYC alter the tumor immune microenvironment to facilitate intravasation and metastasis.

While the specific molecular mediator of TAM recruitment is unknown, we speculate that MYC acts indirectly. Consistent with this, we identified several chemokines/cytokines that were upregulated in Met^High^ tumors and could also be induced by Myc overexpression. Thus, we hypothesize that multiple secreted factors, as opposed to a single factor, act in concert to drive the MYC-associated increase in pro-invasive macrophages.

Most patients with PDAC develop metastases. Our data show that even within this population, the extent of metastatic disease varies widely between patients and impacts survival. While many steps are required for tumor cells to metastasize, our data indicate that bloodstream invasion may be a rate limiting event for metastasis in PDAC. Our work further suggests that in addition to its well-documented cell autonomous role in tumor growth, MYC acts non-cell autonomously to promote metastasis. Given that MYC family members are focally amplified in 28% of human cancer (*73*), these results have broad implications for metastasis in tumor types other than PDAC.

## Acknowledgements

We are grateful for helpful advice from Gregory Beatty, Chi Dang, David DeNardo, and all members of the Stanger Laboratory. This work was supported by grants from the NIH (CA229803 to B.Z.S., K08DK109292 to R.M., F31CA236269 to R.N.), CPRIT (RR190029 to R.M.), AGA (Caroline Craig and Damian Augustyn Award in Digestive Cancer to R.M.). T.B. is supported by the William C. and Joyce C. O’Neil Charitable Trust, Memorial Sloan Kettering Single Cell Sequencing Initiative. F.N. is supported by the Ontario Institute for Cancer Research (OICR), the Canadian Institutes of Health Research (no. 388785), Cancer Research Society (no. 23383), and the Gattuso-Slaight Personalized Cancer Medicine Fund from Princess Margaret Cancer Centre. Additional support was provided by the Abramson Family Cancer Research Institute, the Abramson Cancer Center, and the NIH/Penn Center for Molecular Studies in Digestive and Liver Diseases.

## Materials and Methods

### Mouse models

All experiments were performed in accordance with the National Institutes of Health policies on the use of laboratory animals and approved by the Institutional Animal Care and Use Committee of the University of Pennsylvania. KPCX mice were generated through a series of backcrosses as previously described(*35*). The Rosa^Confetti^ (“X”) reporter allele was introduced into mutant strains bearing Pdx1CreER (“C”), Kras^G12D^ ^(^“K”), and p53^fl/+^ (“P”) alleles to obtain Pdx1CreER; Kras^G12D^; p53^fl/+^; Rosa^Confetti^ (“KPCX”) mice(*35*). For most experiments, animals were heterozygous for the confetti reporter and also contained the Rosa^YFP^ in lieu of the second confetti allele to generate “KPCXY” mice. The Yfp reporter was introduced to enable fluorescent lineage labeling of tumors that undergo a “no-color” recombination event in the confetti reporter as previously described(*74*). To induce recombination, a suspension of TAM (MP Biomedicals) in corn oil (Sigma Aldrich) was administered to pups via lactation following oral gavage of the mother with 6 mg of the drug on post-natal day 0, 1, 2. On average, tumor bearing KPCXY mice were 14-16 weeks of age at time of sacrifice.

### Multicolor image analysis

Pancreatic tumors and organs form tumor bearing KCPXY mice were isolated and analyzed by fluorescent stereomicroscopy using the Lecia M216FA fluorescent microscope with CFP, YFP, and dsRED filters (Chroma). As previously described(*35*), distinct colorimetric tumor clones in the primary tumor mass are defined as an anatomically contiguous region of monochromatic cells that share distinct borders with adjacent clones of a different color (Fig. 1d, Extended Data Fig. 1c). In order to accurately quantify the contribution of a different colorimetric tumors to metastasis we used the following criteria to identify KPCXY mice suitable for analysis: (1) presence of at least one metastatic lesion to the liver and/or lung, (2) two or more tumors present, (3) each metastatic primary tumor carries a unique fluorescent color, and (4) metastatic lesions can be linked to specific tumor based on a shared unique fluorescent lineage label. Using these criteria, we identified a panel of 30 mice with a total of 85 tumors. Metastasis were quantified by fluorescent stereomicroscopy. Tumor size was determined using ImageJ to measure the largest circumference of each fluorescent tumor.

### Murine tumor and metastasis sample acquisition

Pancreatic tumors and associated liver and lung tissues were isolated from tumor bearing KPCXY mice. Under fluorescent stereomicroscopy, individual colored tumors were identified and biopsied using a 6mm punch biopsy. Initial biopsy was placed in 750ml of RNAlater (Sigma Aldrich) for downstream DNA/RNA isolation. Subsequent biopsies were submitted for cell line generation and histology. In tumor where sufficient tissue was available, additional biopsies were taken from anatomically distinct regions of the tumor to obtain subclonal biopsies for genomic analysis (Fig 2a). From 7 KPCXY mice, we obtained biopsies from 20 tumors, 8 of which were amenable to additional sub-regional biopsies. Paired primary tumor and metastasis in each mouse were identified by shared fluorescent lineage labels. Metastasis were harvested by microdissection under fluorescent stereomicroscopy and placed in 500ml of RNAlater for DNA/RNA isolation or cell line generation. Remainder of tissue was embedded for histology. In total, 56 metastases were isolated for genomic analysis. While individual liver metastases were of sufficient size for microdissection, lung lesions were microscopic and could not be readily isolated for molecular analysis.

### Immunofluorescence and histological analysis

Tissue samples were fixed in 4% paraformaldehyde (EMS) at room temperature for 45 minutes followed by an overnight incubation in 30% weight/volume sucrose solution (Sigma Aldrich). Samples were then embedded in O.C.T. (Tissue-Tek) and frozen on dry ice. Staining was performed on 10 μm sections by first blocking with 5% donkey serum and 0.1% Tween-20 for 1 h followed by overnight incubation with primary antibody diluted in blocking buffer in a humidified chamber. Sections were washed three times in PBS containing 0.1% Tween-20. For Immunofluorescence (IF) staining slides were then incubated with DAPI (Life Technologies, 1:1000), and Alexa flour conjugated antibodies (Jackson ImmunoResearch). For immunohistochemistry (IHC), slides were first incubated with biotinylated secondary antibodies (Jackson ImmunoResearch) followed by development using the ABC HRP and DAB kits per manufactures protocol (Vectorlabs). Primary antibodies used were as follows: rat anti-Ki67 (eBioscience, 14-5698-82), rabbit anti-c-Myc [Y69] (Abcam, Ab32072), rabbit anti-CD3 (Invitrogen, PA1-29547), rabbit anti-F4/80 (Novus, NBP2-12506), rat anti-neutrophil (ABCAM, NIMP_R14), and Rabbit Anti-Arg1 (Cell Signaling, 93668).

### Tumor digestion and cell lines

Pancreatic tumors were dissociated into single-cell suspensions through mechanical separation and enzymatic digestion as previously described(*75*). Murine PDAC cell lines 471_Met^High^_1, 832_Met^High^_1, 836_Met^High^_1, 850_Met^High^_4, 852_Met^High^_1, 853 Met^High^_1, 471_Met^Low^_2, 832_Met^Low^_2, 842_Met^Low^_2, 850_Met^Low^_1, 852_Met^Low^_2 were derived from KPCXY mice primary tumors that were also evaluated by SCNA and RNAseq (Fig. 3c and Extended Data Fig. 2). Murine cell lines were cultured in Dulbeccos’ Modified Eagle Medium/F12 medium supplemented with 5 mg/mL D-glucose (Invitrogen), 0.1 mg/mL soybean trypsin inhibitor type I (Invitrogen), 5 mL/L insulin-transferrin-selenium (ITS Premix; BD Biosciences), 25 μg/mL bovine pituitary extract (Gemini Bio-Products), 5 nmol/L 3,3’,5-triiodo-L-thyronine (Sigma), 1 μmol/L dexamethasone (Sigma), 100 ng/mL cholera toxin (Sigma), 10 mmol/L nicotinamide (Sigma), 5% Nu-serum IV culture supplement (ThermoFisher Scientific), and antibiotics (gentamicin 150 μg/mL, Gibco; amphotericin B 0.25 μg/mL, Invitrogen) at 37°C, 5% CO2, 21% O2 and 100% humidity. Cell lines were maintained and passaged according to ATCC recommended procedures and regularly tested for mycoplasma using MycoAlert Mycoplasma Detection Kit (Lonza).

### Orthotopic, subcutaneous, and Circulating Tumor cell analysis

Mice were anaesthetized using isoflurane. For orthotopic injections, NOD.SCID mice were shaved and their abdomen was sterilized using iodine followed by ethanol. An incision was made over the left upper quadrant of the abdomen, and the pancreas was exteriorized. 1.0×10^4^ tumor cells in 50 μl sterile PBS were injected into the pancreas via a 27-gauge 5/8’’ insulin needle. The pancreas was returned to the peritoneal cavity, and the peritoneum and overlying skin were closed with 4-0 coated Vicryl violet FS-2 sutures (Ethicon). Tumors, lungs, and livers were harvested, weighed, measured, and imaged 4-6 weeks following implantation. Blood was drawn from tumor-bearing animals via cardiac puncture with a 1 mL insulin syringe coated with 0.5M EDTA pH 8.0 (Gibco) to prevent coagulation and was immediately placed in a 150mm plate containing 5-10ml RBC lysis buffer (BD Biosciences). After 10 min of lysis at room temperature, PBS was added to the plate and CTCs were directly visualized on a fluorescent microscope and counted. Subcutaneous injections were performed under anesthesia with 1×10^5^ tumor cells in NOD.SCID mice. Tumors were harvested 3-4 weeks after injection.

### Lung metastasis colonization assay

1.0×10^5^ tumor cells were injected via tail vein into NOD.SCID mice. After 21 days, lungs were fluorescently imaged. The tail vein injection experiment was performed in one experiment, 3 mice per group (Met^High^ and Met^Low^) for analysis. Total metastasis counts were quantified using fluorescent stereomicroscopy.

### Immune flow cytometry analysis

Subcutaneous tumors following 20-30 days of implantation were chopped into small pieces and digested in collagenase (1 mg/mL in DMEM; Sigma-Aldrich) at 37°C for 30 minutes and filtered through a 70-μM cell strainer. Single-cell suspensions were stained with LIVE/DEAD Fixable Aqua (Thermo Scientific L-34966) for 10 minutes on ice, followed by antibodies on ice for 30 minutes and washed twice with PBS with 5% FBS for flow cytometric analysis. No intracellular staining is needed for this analysis. Cells were then analyzed by flow cytometry using BD FACS (BD Biosciences) and FlowJo software (Treestar). Antibodies used for the analysis: anti-CD11b PerCP-Cy5.5 (M1/70; BD 550993), anti-F4/80 APC/Cy7 (BM8; Biolegend 123118), anti-I-A/I-E (MHCII) PE/Cy7 (M5/114.15.2; Biolegend 107630), anti-Ly-6C BV570 (HK1.4; Biolegend 128030), anti-Ly-6G V450 (1A8; BD 560603) anti-CD45 AF700 (30-F11; Biolegend 103128), anti-CD206 APC (C068C2; Biolegend 141708), anti-CD115 (CSF-1R) AF488 (AFS98; Biolegend 135512). Gating Strategies for immune cells are as follows: Macrophages – CD45+CD11b+F4/80+, CSF-1R+ Myeloid cells – CD45+CD11b+CSF1R+, Neutrophils – CD45+CD11b+F4/80-Ly6G+. The flow analysis of immune infiltration of Met^Low^ and Met^High^ tumors were performed in two experiments with 4-5 mice per tumor cell clone.

### Bone marrow derived Macrophage isolation

Bone marrow immune cells were isolated as shown previously(*76*). 2×10^6^ isolated immune cells were plated in a six well dish in IMDM (Gibco, 12440053) supplemented with 10% fetal bovine serum (Gibco), 1% L-glutamine (Corning, MT25005CI), 1% Non-essential Amino Acids (Corning, 11140076), 1% Sodium pyruvate (Gibco, 11360-070), 0.001% 2-mercaptoethanol (Gibco, 21985023), 1% Penicillin-Streptomycin (Gibco, 15140163), 20ng/ml recombinant murine M-CSF (PeproTech, 315-02) at 37°C, 5% CO2, 21% O2 and 100% humidity. Bone marrow derived macrophages (BMDMs) were differentiated for 7 days and used by gentle scraping before 10 days

### Macrophage Transwell migration assay

Macrophage invasion was assessed using a 12 well transwell chamber with 8 mm filter inserts (Corning). Growth factor reduced Matrigel (Corning 356231) was diluted 1:1 in PBS and plated onto the transwell. Tumor cells were plated to the lower chamber 24 hours before addition of bone marrow derived macrophages (BMDMs). 100,000 BMDMs were plated in the transwell. After 24 hours, the non-migrated BMDMs and Matrigel were gently removed with a swab. Cells in the lower surface (migrated BMDMs) of the membrane were fixed in 4% paraformaldehyde (PFA) for 15 mins. DAPI (Invitrogen D21490) was added in PBS to the transwells. The membranes were imaged and number of macrophages counted in 4 random fields. The experiment was performed in triplicates and was repeated twice.

### Macrophage Depletion in vivo

10,000 tumor cells were orthotopically injected into the pancreas of NOD.SCID mice. Treatments started 10 days after implantation. GW2580, CSF-1R inhibitor (AdooQ Bioscience, A11959-200) was dissolved in 0.5% hydroxypropyl methyl cellulose and 0.1% Tween and dosed 3 times a week at 160mg/kg by oral gavage. 200ul of Clodronate or control liposomes (Liposoma, CP-025-025) was given one time a week by intraperitoneal injection. Blood from tail snips was used to analyze depletion. Experimental and control mice were euthanized 14 days after treatment start and analyzed for metastasis and immune cells by flow cytometry. Control and experimental groups were run in triplicate.

### Molecular cloning, Lentivirus generation and transduction

Murine *Myc* was cloned into pCDH-EF1-FHC (Addgene plasmid # 64874). The vectors were transfected into 293T cells using PEI reagent and packaged into lentivirus for transduction. Lentivirus was collected 48 hours after transfection and 0.45um filtered for usage. Tumor cells were transduced with packaged virus with 4ug/ml polybrene for 48 hours. Cells were selected with puromycin for 10 days. Overexpression efficiencies were assessed by gene-specific quantitative PCR analysis.

### Immunoblotting

Cells were washed with cold PBS and lysed in RIPA buffer on ice. Lysis was spun down at max speed and supernatant was collected. Protein was separated by SDS-PAGE, transferred to PVDF membrane, blocked in 5% non-fat milk in PBS plus 0.1% Tween-20, probed with primary antibodies, and detected with horseradish peroxidase- conjugated secondary antibodies (Jackson ImmunoResearch). Primary antibodies used include: anti-Myc [Y69] (Abcam, ab32072); anti-GAPDH (CST, 2118S)

### Real-time PCR

RNA was prepared from cultured tumor cells using RNeasy Mini Kit (Qiagen). cDNA was generated using High-capacity cDNA Reverse Transcription Kit in a 20 μl reaction volume and diluted 1:10 for qPCR analysis (Life Technologies). qPCR analysis was performed using 1 μl diluted cDNA with biological (2-3) and technical replicates (2-3) using SsoAdvanced SYBR reagent (Bio-Rad) and Bio-Rad qPCR platform, and results were normalized to the expression of Gapdh using the Bio-Rad software. Primer sequences utilized for qPCR were: mGapdh F- atgttccagtatgactccactcacg, mGapdh R- gaagacaccagtagactccacaca, mMyc F- gcatgaggagacaccgccca, mMyc R- ggtttgcctcttctccacaga

### DNA and RNA isolation and purification

Tumor and metastasis biopsy samples were stored in RNAlater (Sigma Aldrich) prior to processing. Bulk tissues were homogenized in triazol (ThermoFisher) and followed by phase separation with 0.2ml chloroform per 1ml of Triazol. Aqueous phase was isolated and RNA extracted with RNAeasy Mini Kit (Quiagen) per manufactures protocol. To extract DNA, the remainder of the solution following removal of aqueous phase was mixed with 300ul of Back extraction buffer (4M Guanidine Thiocyanate, 50mM Sodium Citrate dihydrate, 1M Tris free base, and molecular grade water. All reagents from Sigma Aldrich) per 1ml of triazol. Solution was incubated on a shaker for 10 min followed by centrifugation for 30 minutes at 12000xg at room temperature. Aqueous solution was isolated and 0.4ml of Isopropanol was added followed by centrifugation for 15 minutes at 12000xg at 4C. The DNA pellet was then washed with 70% ethanol.

---

**Supplementary Fig. 1:**
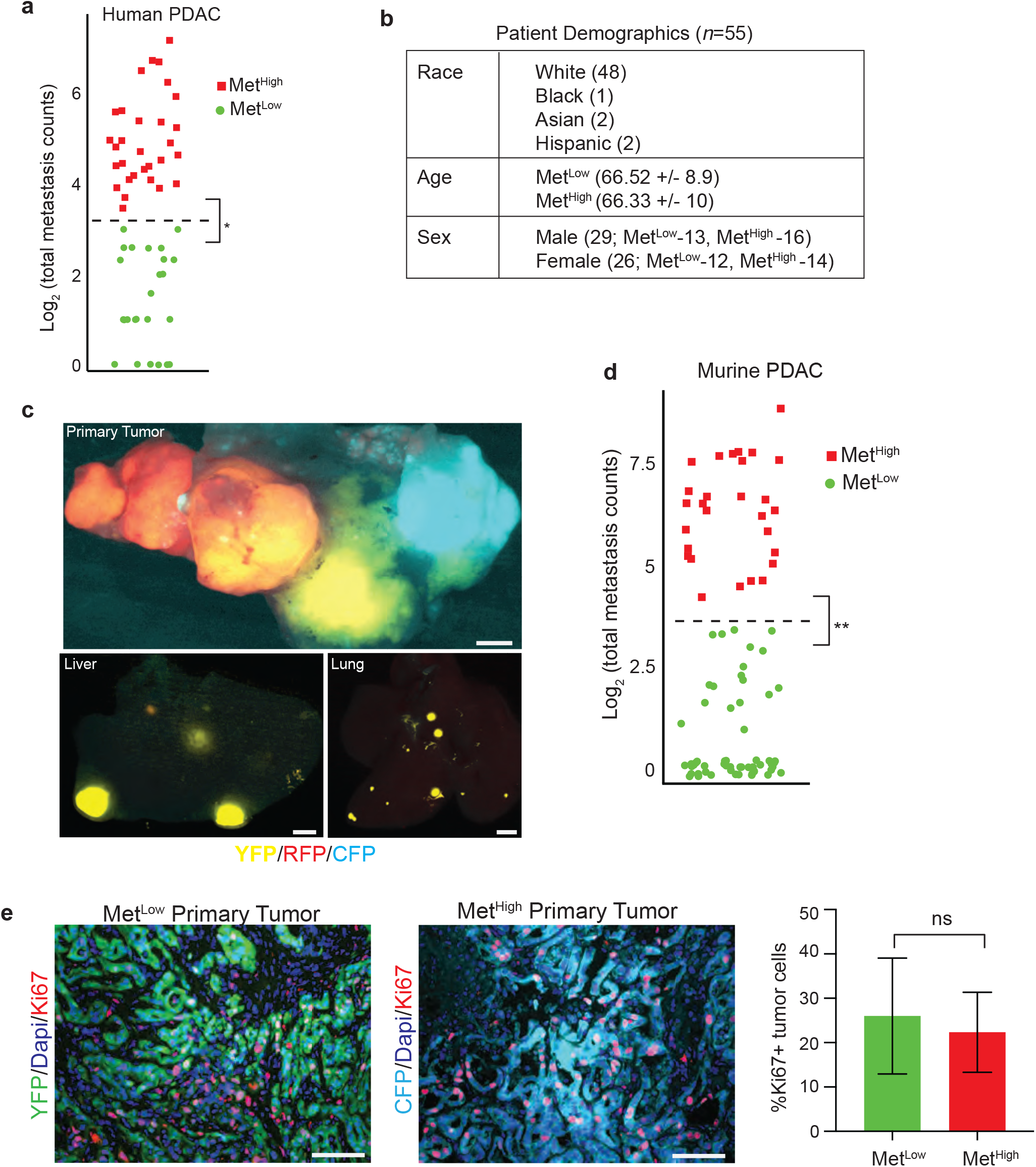
Classifying metastatic burden in human and murine advanced PDA. **(a)** K-means clustering of human PDAC metastasis counts demonstrating two distinct clusters with Met^Low^ ≤10 and Met^High^ >10 total (liver and lung) metastasis. n=55 patients. **(b)** Demographics (age of diagnosis, sex, and race) of the patients analyzed in Fig. 1a-d and Extended Data Fig. 1a. **(c)** Top: Representative stereomicroscopic fluorescent image showing multiple primary tumors (RFP+, YFP+, and CFP+) in the pancreas with matched metastases in the liver and lung. Liver and lung metastases are derived primarily from the YFP+ tumor. **(d)** K-means clustering of murine PDAC metastasis counts demonstrating two distinct clusters that are defined as having high or low metastatic burden. Met^Low^ ≤10 and Met^High^ >10 total (liver and lung) metastasis, n=85 tumor clones. **(e)** Ki67 staining in primary Met^Low^ and Met^High^ tumors with representative IF images (left) and counts (right). Data from n=3 Met^Low^ and n=3 Met^High^ primary tumors and 4-5 random fields of view. Statistical analysis by students unpaired t-test with significance indicated (ns, not significant). Error bars indicating SEM. Scale bar 1mm for Extended Data Fig. 1c and 50um for Extended Data Fig. 1e.

**Supplementary Fig. 2.**
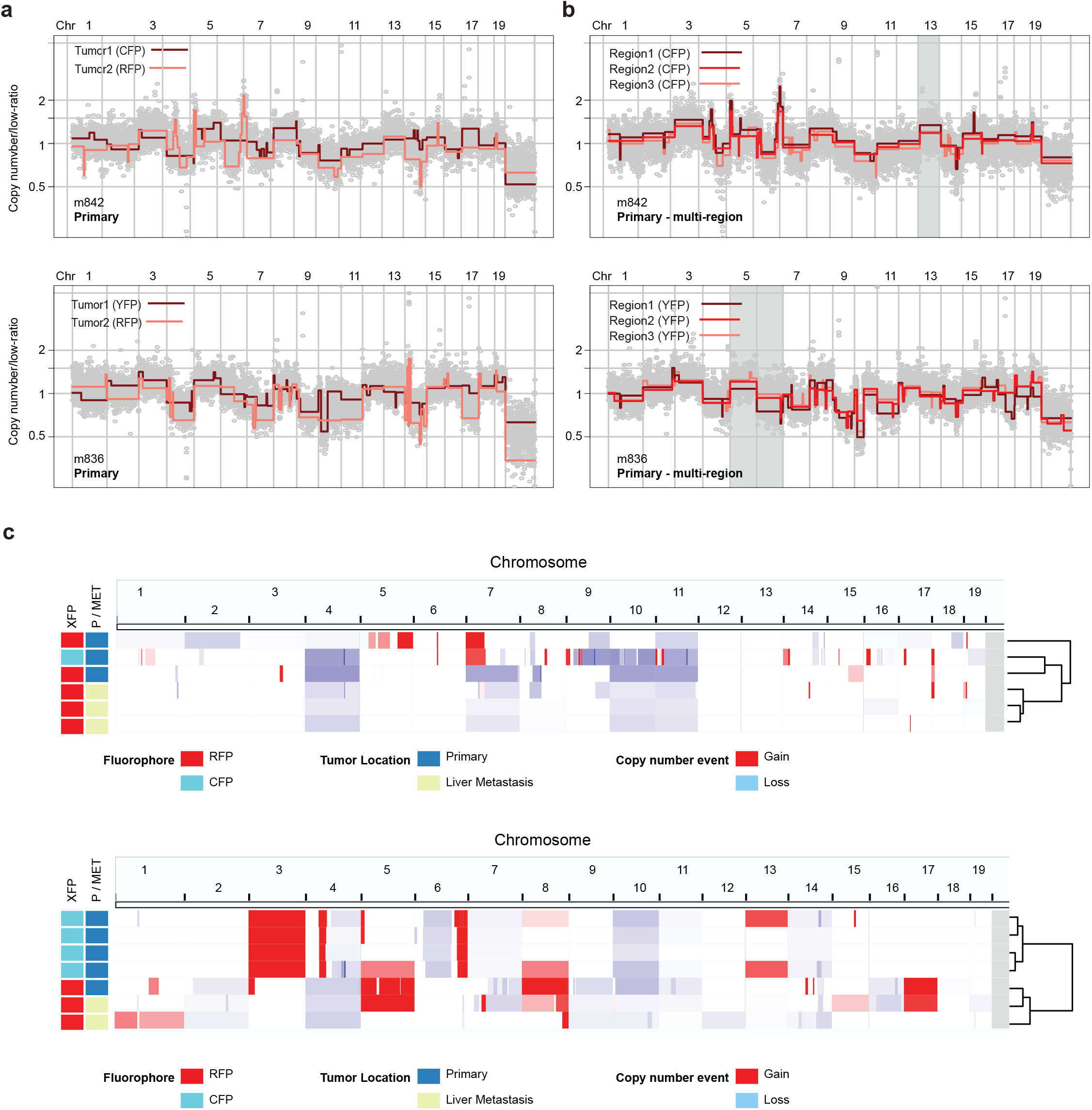
Copy number alteration analysis reveals lineage relationships and genetic heterogeneity of fluorescently labeled primary PDACs and their matching metastasis. **(a)** Overlay genome-wide copy number profiles of different fluorescently labeled (CFP and RFP) primary tumors from mouse 842 (m842 - top panel) and mouse 836 (m836 - bottom panel) illustrating differing rearrangement profiles. **(b)** Overlay genome-wide copy number profiles of tumors where multi-region sampling was performed. Top panel illustrates three CFP sub-samples from CFP tumor mass from m842. Bottom panel illustrates three YFP sub-samples from a YFP tumor mass from m836. **(c)** Genome-wide heatmap with hierarchal clustering based on copy number alterations of matched primary and metastatic samples profiled from m471 (top panel) and m842 (bottom panel). Color codes for Flourescence label, primary/metastatic designation, and nature of copy number alteration are provided below.

**Supplementary Fig. 3:**
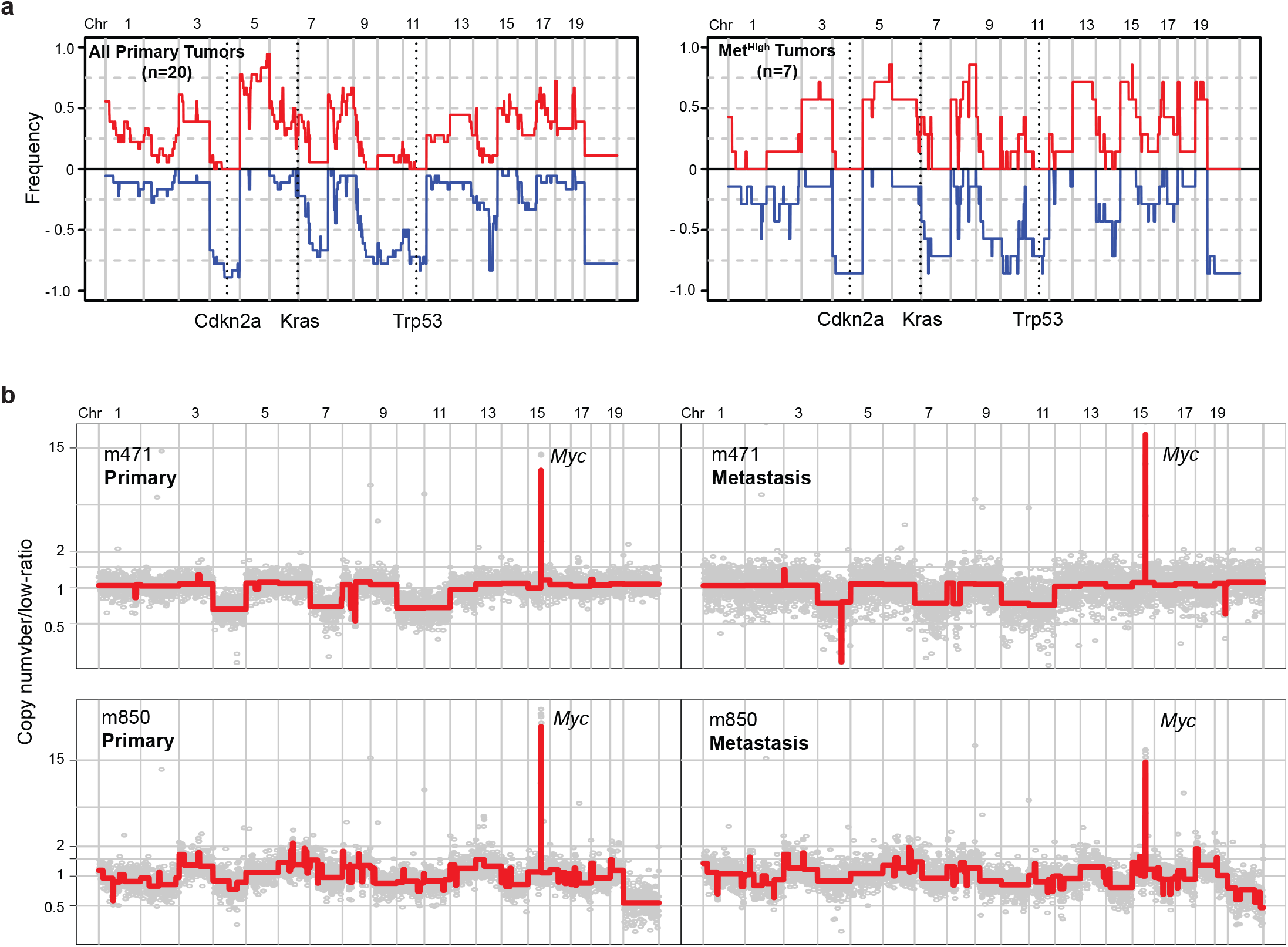
Global patterns of large scale copy number alterations do not differ between Met^High^ and Met^Low^ clones, while Myc amplifications are maintained in metastatic lesions. **(a)** Genome-wide frequency plot illustration of large scale copy number events found in all primary tumors (left panel) and in Met^High^ (right panel) primary tumors. Alterations in Cdkn2a, Kras, and Trp53 are noted on the plots. **(b)** Genome-wide copy number profiles of matching Met^High^ primary tumors (left panels) and paired liver metastases (right panels) from different examined mice. Myc amplifications are noted on profiles.

**Supplementary Fig. 4:**
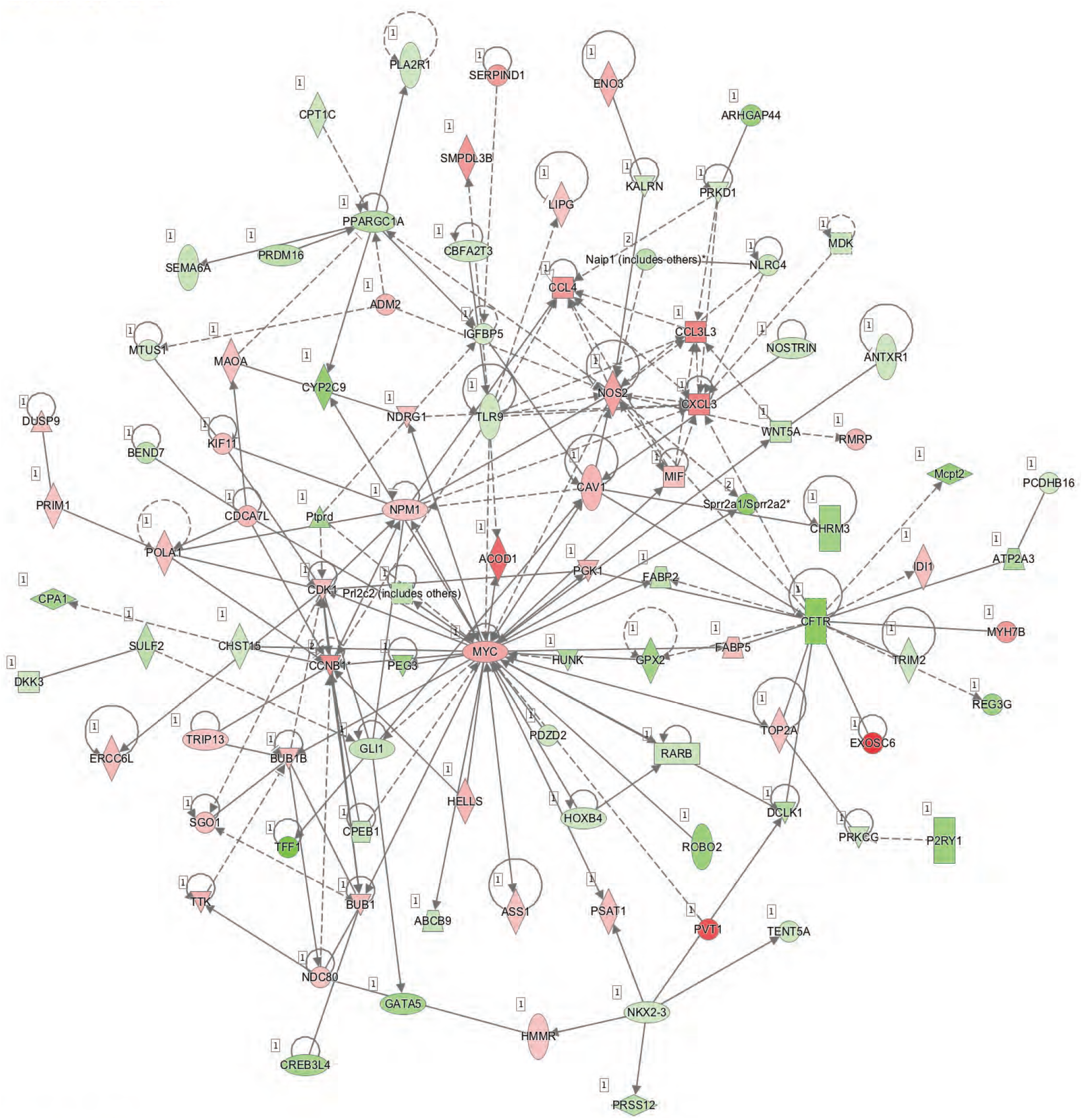
Ingenuity pathway analysis of the Met^High^ transcriptome.

**Supplementary Fig. 5:**
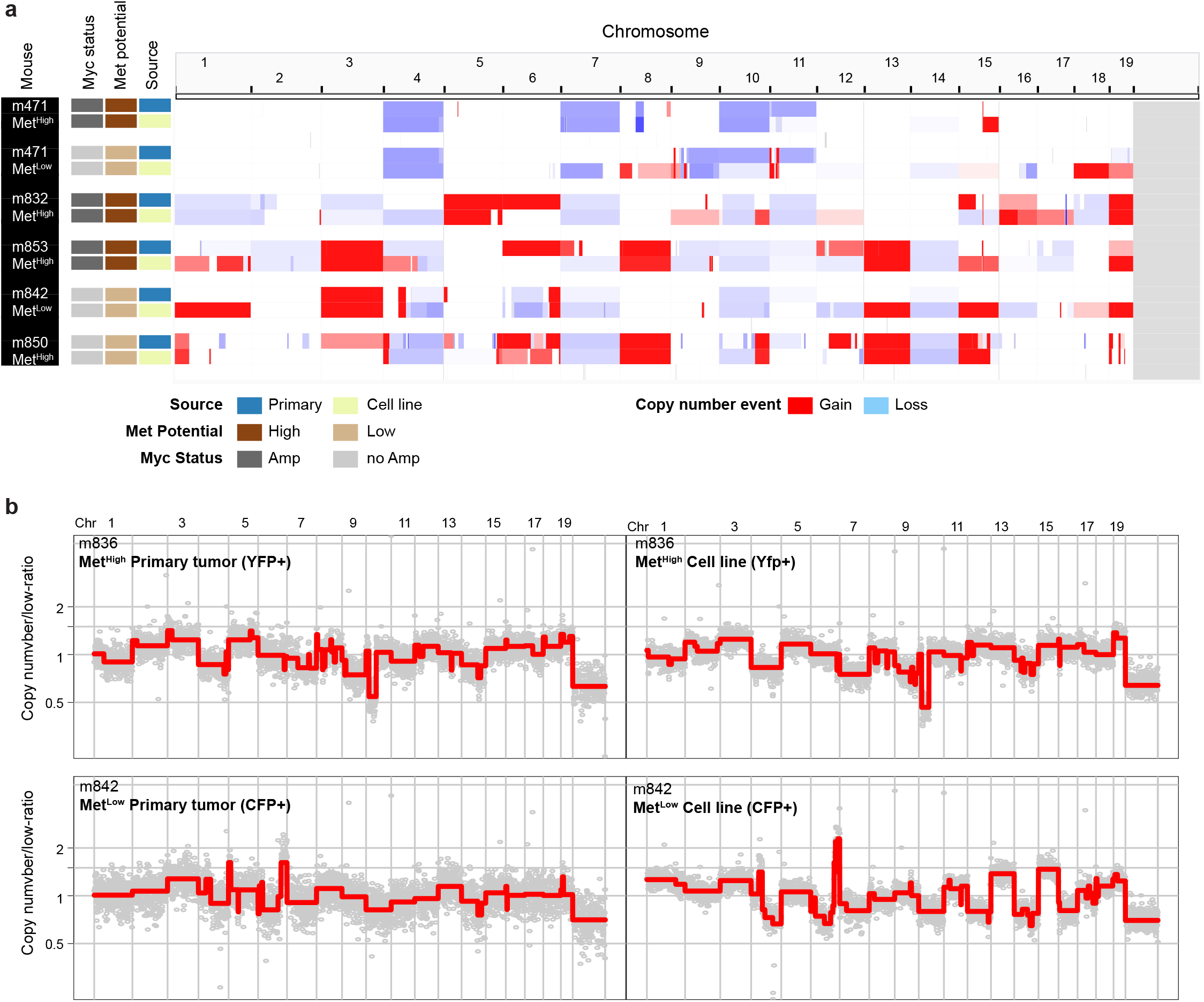
Established Met^High^ / Met^Low^ cell lines maintain their overall genomic profile and do not select for Myc amplification during in vitro culture. **(a)** Schematic representation of genome wide copy number profile of matching primary tumor and cell lines samples illustrating the maintenance of their overall copy number pattern. (b) Representative genome-wide copy number profiles of matching primary tumors and cell line samples for a Met^High^ tumor from m836 (YFP+) and Met^Low^ tumor from m842 (CFP+), top and bottom panels respectively.

**Supplementary Fig. 6:**
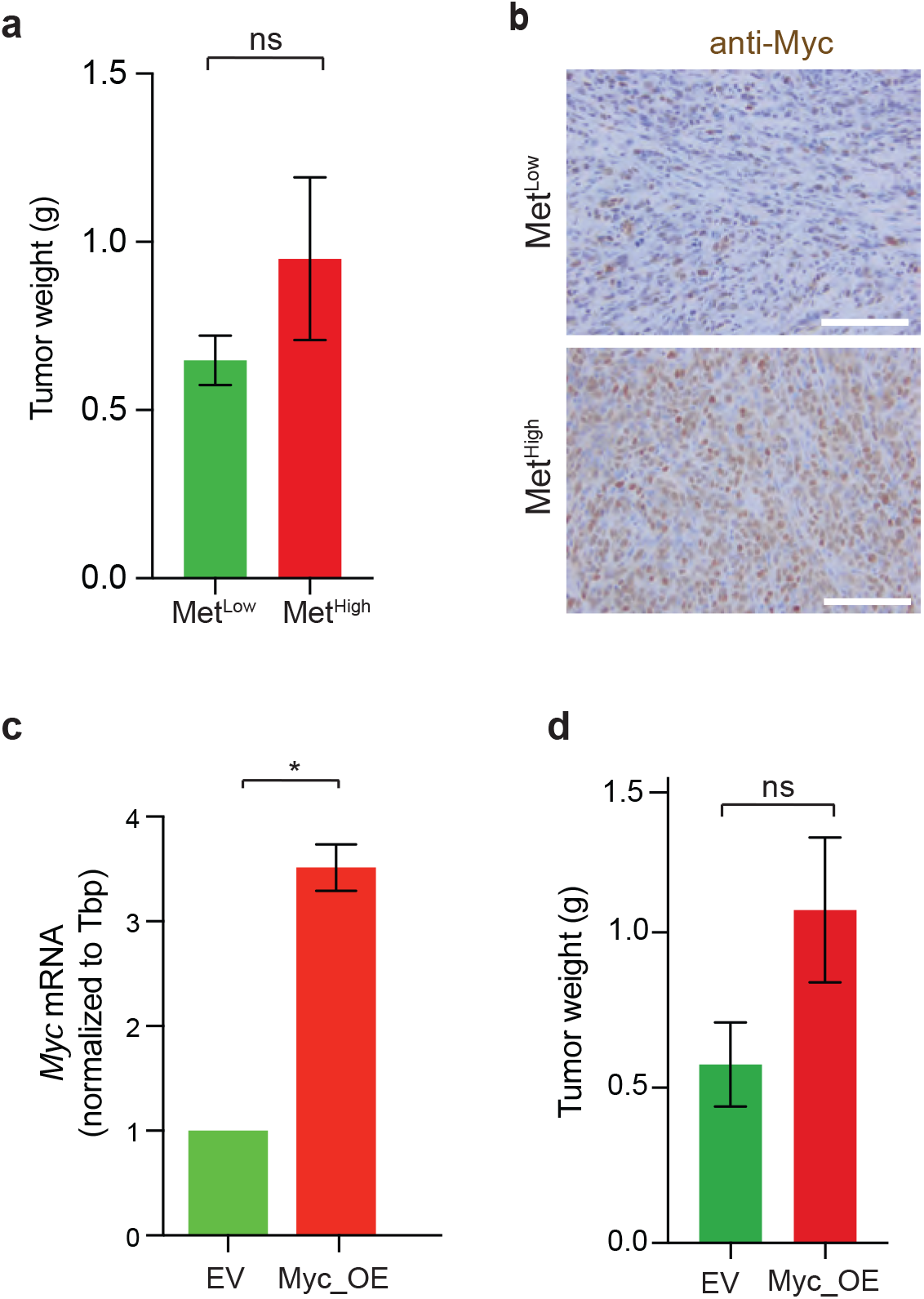
Tumor weight and Myc expression in orthotopic primary tumors primary and Myc overexpressing cell lines. **(a)** Tumor weight in grams of Met^Low^ and Met^High^ cell line derived orthotopic tumors in Figure 4c-d. Weight from n= 4 Met^Low^ and 5 Met^High^ cell lines. **(b)** Representative IHC staining of Met^Low^ and Met^High^ orthotopic tumors from Figure 4c-d. **(c)** Myc gene expression in Met^Low^_Myc-OE relative to Met^Low^_EV cell lines in Figure 4e-f. **(d)** Tumor weight in grams of Met^Low^_EV and Met^Low^_Myc-OE cell line derived orthotopic tumors in Figure 4e-f. Statistical analysis by Student’s unpaired t-test with significance indicated (*, *p*=0.007; ns, not significant). Error bar indictes SEM (panels a,c,d). Scale bar 50mm (panel b).

**Supplementary Fig. 7:**
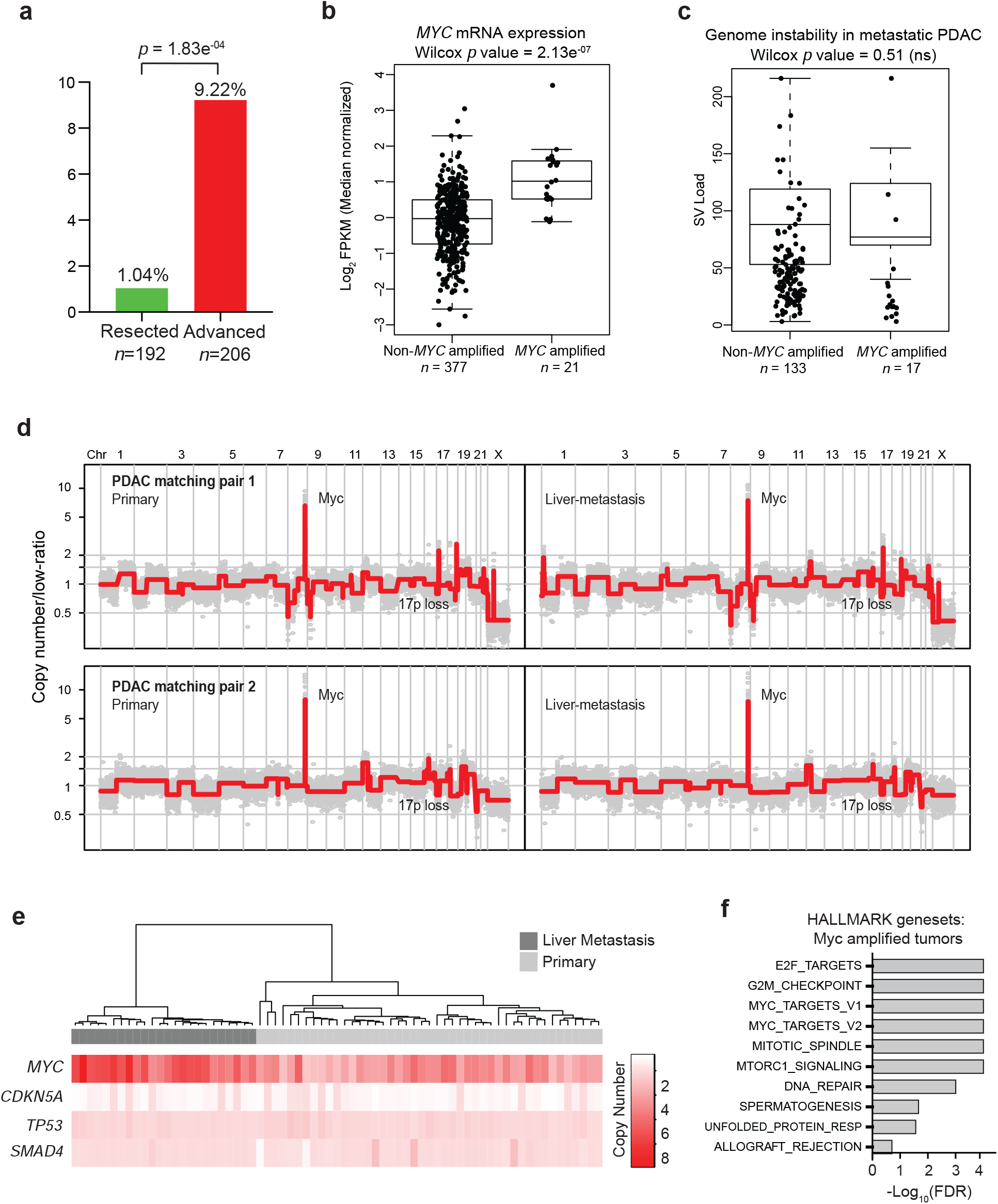
*MYC* amplifications are enriched in metastatic human PDA. **(a)** Bar graph showing the relative frequencies of MYC amplifications in resected and advanced PDAC tumors from the COMPASS cohort. **(b)** *MYC* expression in non-MYC amplified and MYC amplified tumors in the COMPASS cohort. **(c)** Measurement of total structural variant burden in non-MYC amplified and MYC amplified tumors in the COMPASS cohort. **(d)** Genome-wide relative copy number profiles of two patients with matched primary PDAC (left) and liver metastasis (right). **(e)** Heatmap depiction of Myc, CDKN5A, TP53, and SMAD4 copy number alteration in cancer single cells sequenced from a matched primary PDAC and its liver metastasis depicted in Fig. 6e. Color codes indicate absolute copy number in single-cells. Top bar plot depicts tissue site from which single-cells were retrieved. (f) Gene set enrichment analysis of MYC amplified tumors (compared to non-amplified tumors) in the COMPASS cohort.

**Supplementary Table 1.** Description of mice and murine tumor clones analyzed.

| Mouse ID | Total # of fluorescent tumor clones | #Met<sup>Low</sup> clones | #Met<sup>High</sup> clones | Clone ID and color | #sub-clonal biopsies analyzed by SCNA | Met<sup>High</sup> Liver metastasis counts | Met<sup>High</sup> Lung metastasis counts |
| --- | --- | --- | --- | --- | --- | --- | --- |
| 471 | 3 | 2 | 1 | 471_Met<sup>High</sup>_1: RFP<br>471_Met<sup>Low</sup>_2: RFP<br>471_Met<sup>Low</sup>_3: CFP | 471_Met<sup>High</sup>_1: 2<br>471_Met<sup>Low</sup>_2: 1<br>471_Met<sup>Low</sup>_3: 1 | 80 | 1 |
| 832 | 2 | 1 | 1 | 832_Met<sup>Low</sup>_1: YFP<br>832_Met<sup>High</sup>_2: CFP | 832_Met<sup>Low</sup>_1: 2<br>832_Met<sup>High</sup>_2: 4 | 45 | 0 |
| 836 | 3 | 2 | 1 | 836_Met<sup>High</sup>_1: YFP<br>836_Met<sup>Low</sup>_2: RFP<br>836_Met<sup>Low</sup>_3: RFP | 836_Met<sup>High</sup>_1: 3<br>836_Met<sup>Low</sup>_2: 1<br>836_Met<sup>Low</sup>_3: 1 | 49 | 130 |
| 842 | 4 | 3 | 1 | 842_Met<sup>Low</sup>_1: CFP<br>842_Met<sup>High</sup>_2: RFP<br>842_Met<sup>Low</sup>_3: RFP<br>842_Met<sup>Low</sup>_4: YFP | 842_Met<sup>Low</sup>_1: 4<br>842_Met<sup>High</sup>_2: 1<br>842_Met<sup>Low</sup>_3: 1<br>842_Met<sup>Low</sup>_4: 2 | 40 | 70 |
| 850 | 4 | 3 | 1 | 850_Met<sup>Low</sup>_1: YFP<br>850_Met<sup>Low</sup>_2: YFP<br>850_Met<sup>Low</sup>_3: CFP<br>850_Met<sup>High</sup>_4: RFP | 850_Met<sup>Low</sup>_1: 1<br>850_Met<sup>Low</sup>_2: 1<br>850_Met<sup>Low</sup>_3: 1<br>850_Met<sup>High</sup>_4: 3 | 53 | 28 |
| 852 | 2 | 1 | 1 | 852_Met<sup>High</sup>_1: YFP<br>852_Met<sup>Low</sup>_2: RFP | 852_Met<sup>High</sup>_1: 3<br>852_Met<sup>Low</sup>_2: 1 | 63 | 52 |
| 853 | 2 | 1 | 1 | 853_Met<sup>High</sup>_1: YFP<br>853_Met<sup>Low</sup>_2: RFP | 853_Met<sup>High</sup>_1: 2<br>853_Met<sup>Low</sup>_2: 1 | 250 | 200 |
| Total | 20 | 13 | 7 |  |  |  |  |

Table shows mouse ID, number of Met<sup>Low</sup> and Met<sup>High</sup> clones from each mouse, number of sub-clonal biopsies analyzed by SCNA analyzed for each primary tumor clone, and metastasis counts to the liver and lung.

## Supplementary Methods

### Analysis of RNA-seq, Differential gene expression, GSEA, and molecular subtype

RNA sequencing was performed on bulk tumor and metastasis samples from 7 KPCX mice resulting in 66 samples for analysis (Primary tumor with subregional biopsies and metastasis). RNA purity and integrity were verified on the Agilent Tapestation prior to library construction followed by paired-end 75bp sequencing on an Illumina HiSeq 4000 high-throughput sequencer. Alignment of fastq files was performed with STAR aligner v2.5.2b using mm10 as the reference genome(*77*). Gene level expression data in terms of expected counts and FPKM was obtained using RSEM v1.2.28(*78*). Low expressing genes were removed using cutoff of 100 for count and 10 for FKPM. For primary tumor clones where subregional biopsies were taken, count data for the tumor was obtained by merging expression data of the subclone count data using the mean value. This resulted in a total of 54 samples for downstream analysis (Met^Low^ = 13, Met^High^ = 7, and Metastasis = 34). For differential expression analysis, count data was normalized using the voom function in the limma R-package followed by batch correction using the ComBat R-package(*79, 80*). Then limma was used to perform differential expression between Met^High^, Met^Low^, and metastasis. Boxplots of log2 FPKM values for genes were generated using the ggplot2 R-package. To generate volcano plots, differential expression data comparing Met^Low^ and Met^High^ clones was plotted using ggplot2 with log base 2-fold change from Met^Low^ compared to Met^High^ tumors of each gene was plotted on the x-axis and the adjusted p-values plotted on the Y axis. Genes with adjusted p-values less than 0.01 and absolute log2 fold change >1 were highlighted. Differentially expressed genes were used as input for GSEA MSigDB geneset enrichment analysis(*81*). Transcription factor enrichment on all differentially expressed genes was performed using the Metacore software package (https://clarivate.com/products/metacore/, Clarivate Analytics, London, UK). Network analysis was performed on all differentially expressed genes using ingenuity pathway analysis software (www.ingenuity.com, Ingenuity Systems Inc., Redwood City, CA). Molecular subtype classification using the Bailey, Moffitt, and Collision(*18, 19, 44*) signature was performed on each sample by subtracting the sum of normalized expression for genes corresponding to specific classes within a particular molecular signature. We then took the maximum score observed across each class and assigned to the samples. Heatmaps were generated using differentially expressed genes with adjusted p-value <0.05 and absolute LogFC >1.

### Survival analysis of TCGA data

TCGA PAAD expression and patient and sample level clinical data was downloaded from cBioportal (http://www.cbioportal.org/). Samples were filtered to those classified as pancreatic adenocarcinoma and having available expression data (162 of 186 samples). To develop a signature gene list associated with the Met^High^ phenotype, we filtered differentially expressed genes between Met^High^ and Met^Low^ tumors using an adjusted p-value <0.05 and absolute logFC >0.58 resulting in a set of genes that are upregulated or downregulated in the Met^High^ tumors (736 Up and 1036 down). To calculate a signature score for each TCGA PAAD sample, we first z-score normalized the TCGA PAAD expression data and then subtracted the sum of all downregulated gene expression from the sum of all upregulated gene expression values. We divided the signature score into high and low strata using a cutoff score >0. Kaplan-Meier analysis was done to compare survival between the two groups.

### Human Stage IV pancreatic tumor and metastasis imaging analysis

CT scans were obtained from patients with metastatic PDAC undergoing treatment at the University of Pennsylvania (IRB protocol #822028). Patients were filtered to include only those with CT-scan imaging of the abdomen and chest with IV contrast at the time of diagnosis and prior to any treatment and found to have Stage IV disease. In total, 55 patients were included. CT images for each patient were reviewed and metastatic lesions in liver and lung were counted. All metastases were examined in multiple planes to ensure accurate assessment. Tumor area was pulled from the initial radiologist report and measured at the largest diameter. Cut-off for high and low metastasis groups was determined using k-means with n=2 clusters.

### Human pancreatic cancer patient sample acquisition with genomic and RNA-seq analysis

Sample acquisition resulted from patients recruited as part of the International Cancer Genome Consortium (ICGC) Pancreatic Cancer Ductal Adenocarcinoma Canadian sequencing initiative or the COMPASS trial as previously described(*82*). Tissue samples were collected at the University Health Network (Toronto), Sunnybrook Health Sciences Centre (Toronto), Kingston General Hospital (Kingston), McGill University (Montreal), Mayo Clinic (Rochester), or Massachusetts General Hospital (Boston) with patient informed consent and approval from Institutional Review or Research Ethics Boards. Whole Genome Sequencing and RNA Sequencing was performed on fresh frozen tumor tissue samples which were enriched for tumor content by laser capture microdissection (LCM). Whole genome sequencing (WGS) and RNAseq were performed at the Ontario Institute of Cancer Research as described previously(*82*). DNA Read Alignment and MYC Copy Number Variations were performed on paired end whole genome sequencing reads aligned to human reference genome hg19 using BWA 0.6.2(*83*). PCR duplicates were marked with Picard 1.90. Tumor cellularity, ploidy, and copy number segments were derived using an in-house algorithm CELLULOID(*84*). RNA reads were aligned to human reference genome hg38 and to transcriptome Ensemble v84 using STAR v2.5.2a(*77*). Duplicate reads were marked with Picard 1.121. Raw counts were obtained using HTSeq 0.6.1(*85*). Differential gene expression analysis was performed with DESeq2 v.1.14.1(*86*) using default settings. Briefly, RNA HTSeq count data was imported to generate a dispersion estimate and a generalized linear model. Wald statistical test was used to compare gene expression of MYC amplified to non-amplified cases. Gene set enrichment analysis was performed using genes ranked based on the P value and sign of the log2 fold change from differential gene expression analysis. Gene set enrichment analysis was run using GSEA Preranked 4.0.2 with default settings against hallmark gene sets(*81*). Statistical Analyses included pairwise comparisons of quantitative variables performed using Wilcoxon rank sum test.. All tests were two-sided. Analyses were carried out in R 3.3.0.

### Somatic Copy Number Analysis in murine tumors

DNA purified from dissected murine tumors was processed for Illumina library preparation and sequencing using standard protocols. In brief, isolated DNA (between 100-100ng in total) was sonicated on a Covaris instrument. Sonicated DNA was then end-repaired and ligated to TruSeq dual-index library adaptors. Index libraries were subsequently enriched by 10 cycles of PCR amplification followed by pooling and multiplex sequencing targeting a coverage of roughly 2 million reads per sample(*87*). For data processing and copy number inference, sequencing data was processed as previously described with mouse genomic bins computed in a manner similar to human bins(*38*). In brief, sequencing reads were mapped to the mouse reference genome built mm9. Sequencing reads were indexed, sorted, with PCR duplicates removed. Uniquely mapped reads were counted in each bin and normalized for GC content using lowess smoothing. Normalized read count data were then segmented using Circular Binary Segmentation (CBS)(*88*) with the profiles centered around a mean of 1. Chromosomal segments with variance that is above or below the mean where called as gains or deletions, respectively. A threshold of 0.2 was used. For hierarchal clustering and lineage reconstruction, an analysis based on copy number values and alteration breakpoints, in a genome-wide manner, was employed.

### Bulk and single cell analysis of matched primary and metastasis from human tissue

For 20 patients, tissue sections from flash frozen samples were processed for bulk DNA purification using Qiagen DNeasy Blood and Tissue kit. Purified DNA was processed as described above for multiplex sequencing. A coverage of 2 million sequencing reads was similarly targeted. For a single case, matched pancreatic primary and liver metastasis tissue was retrieved and processed for single-nuclei isolation as previously described(*38*). Single-nuclei were sorted based on DNA content form both diploid and polyploid populations of each tissue. Approximately 100 nuclei per tissue sample was amplified using WGA4 kit (Sigma-Aldrich) with the resulting Whole Genome Amplified (WGA) DNA processed for TruSeq Indexed sequencing library preparation as described above. Sequencing data was processed as described above with the exception that a least squares fitting algorithm was used to calculate absolute integer copy number(*87*).

## References

1. N. McGranahan, C. Swanton, Clonal Heterogeneity and Tumor Evolution: Past, Present, and the Future. Cell 168, 613–628 (2017).

2. S. Turajlic et al., Tracking Cancer Evolution Reveals Constrained Routes to Metastases: TRACERx Renal. Cell 173, 581–594 e512 (2018).

3. K. W. Hunter, R. Amin, S. Deasy, N. H. Ha, L. Wakefield, Genetic insights into the morass of metastatic heterogeneity. Nat Rev Cancer 18, 211–223 (2018).

4. G. Gundem et al., The evolutionary history of lethal metastatic prostate cancer. Nature 520, 353–357 (2015).

5. Z. M. Zhao et al., Early and multiple origins of metastatic lineages within primary tumors. Proceedings of the National Academy of Sciences of the United States of America 113, 2140–2145 (2016).

6. C. C. Foster, S. P. Pitroda, R. R. Weichselbaum, Definition, Biology, and History of Oligometastatic and Oligoprogressive Disease. Cancer J 26, 96–99 (2020).

7. S. P. Pitroda, R. R. Weichselbaum, Integrated molecular and clinical staging defines the spectrum of metastatic cancer. Nat Rev Clin Oncol 16, 581–588 (2019).

8. R. R. Weichselbaum, S. Hellman, Oligometastases revisited. Nat Rev Clin Oncol 8, 378–382 (2011).

9. M. P. Deek, P. T. Tran, Oligometastatic and Oligoprogression Disease and Local Therapies in Prostate Cancer. Cancer J 26, 137–143 (2020).

10. R. Phillips et al., Outcomes of Observation vs Stereotactic Ablative Radiation for Oligometastatic Prostate Cancer: The ORIOLE Phase 2 Randomized Clinical Trial. JAMA Oncol 6, 650–659 (2020).

11. A. J. Weickhardt et al., Local ablative therapy of oligoprogressive disease prolongs disease control by tyrosine kinase inhibitors in oncogene-addicted non-small-cell lung cancer. J Thorac Oncol 7, 1807–1814 (2012).

12. D. P. Ryan, T. S. Hong, N. Bardeesy, Pancreatic adenocarcinoma. N Engl J Med 371, 1039–1049 (2014).

13. S. Mueller et al., Evolutionary routes and KRAS dosage define pancreatic cancer phenotypes. Nature 554, 62–68 (2018).

14. M. Chan-Seng-Yue et al., Transcription phenotypes of pancreatic cancer are driven by genomic events during tumor evolution. Nature genetics 52, 231–240 (2020).

15. R. A. Moffitt et al., Virtual microdissection identifies distinct tumor- and stroma-specific subtypes of pancreatic ductal adenocarcinoma. Nature genetics 47, 1168–1178 (2015).

16. P. Bailey et al., Genomic analyses identify molecular subtypes of pancreatic cancer. Nature 531, 47–52 (2016).

17. P. J. Campbell et al., The patterns and dynamics of genomic instability in metastatic pancreatic cancer. Nature 467, 1109–1113 (2010).

18. a. a. d. h. e. Cancer Genome Atlas Research Network. Electronic address, N. Cancer Genome Atlas Research, Integrated Genomic Characterization of Pancreatic Ductal Adenocarcinoma. Cancer cell 32, 185–203 e113 (2017).

19. A. El-Kenawi, K. Hanggi, B. Ruffell, The Immune Microenvironment and Cancer Metastasis. Cold Spring Harb Perspect Med 10, (2020).

20. A. Swierczak, J. W. Pollard, Myeloid Cells in Metastasis. Cold Spring Harb Perspect Med, (2019).

21. T. Kitamura et al., CCL2-induced chemokine cascade promotes breast cancer metastasis by enhancing retention of metastasis-associated macrophages. J Exp Med 212, 1043–1059 (2015).

22. J. Wyckoff et al., A paracrine loop between tumor cells and macrophages is required for tumor cell migration in mammary tumors. Cancer research 64, 7022–7029 (2004).

23. M. D. Wellenstein, K. E. de Visser, Cancer-Cell-Intrinsic Mechanisms Shaping the Tumor Immune Landscape. Immunity 48, 399–416 (2018).

24. S. K. Wculek, I. Malanchi, Neutrophils support lung colonization of metastasis-initiating breast cancer cells. Nature 528, 413–417 (2015).

25. B. M. Szczerba et al., Neutrophils escort circulating tumour cells to enable cell cycle progression. Nature 566, 553–557 (2019).

26. B. Z. Qian et al., CCL2 recruits inflammatory monocytes to facilitate breast-tumour metastasis. Nature 475, 222–225 (2011).

27. J. Park et al., Cancer cells induce metastasis-supporting neutrophil extracellular DNA traps. Sci Transl Med 8, 361ra138 (2016).

28. N. Linde et al., Macrophages orchestrate breast cancer early dissemination and metastasis. Nat Commun 9, 21 (2018).

29. D. G. DeNardo et al., CD4(+) T cells regulate pulmonary metastasis of mammary carcinomas by enhancing protumor properties of macrophages. Cancer cell 16, 91–102 (2009).

30. J. Li et al., Tumor Cell-Intrinsic Factors Underlie Heterogeneity of Immune Cell Infiltration and Response to Immunotherapy. Immunity 49, 178–193 e177 (2018).

31. A. Pommier et al., Unresolved endoplasmic reticulum stress engenders immune-resistant, latent pancreatic cancer metastases. Science 360, (2018).

32. R. Maddipati, B. Z. Stanger, Pancreatic Cancer Metastases Harbor Evidence of Polyclonality. Cancer Discov, (2015).

33. S. Yachida, C. A. Iacobuzio-Donahue, The pathology and genetics of metastatic pancreatic cancer. Arch Pathol Lab Med 133, 413–422 (2009).

34. C. A. Iacobuzio-Donahue et al., DPC4 gene status of the primary carcinoma correlates with patterns of failure in patients with pancreatic cancer. J Clin Oncol 27, 1806–1813 (2009).

35. N. Navin et al., Inferring tumor progression from genomic heterogeneity. Genome Res 20, 68–80 (2010).

36. T. Baslan et al., Genome-wide copy number analysis of single cells. Nature protocols 7, 1024–1041 (2012).

37. D. J. H. Shih et al., Genomic characterization of human brain metastases identifies drivers of metastatic lung adenocarcinoma. Nature genetics 52, 371–377 (2020).

38. R. Klotz et al., Circulating Tumor Cells Exhibit Metastatic Tropism and Reveal Brain Metastasis Drivers. Cancer Discov 10, 86–103 (2020).

39. R. Dhanasekaran et al., MYC and Twist1 cooperate to drive metastasis by eliciting crosstalk between cancer and innate immunity. Elife 9, (2020).

40. O. G. McDonald et al., Epigenomic reprogramming during pancreatic cancer progression links anabolic glucose metabolism to distant metastasis. Nature genetics 49, 367–376 (2017).

41. S. H. Chiou et al., BLIMP1 Induces Transient Metastatic Heterogeneity in Pancreatic Cancer. Cancer Discov 7, 1184–1199 (2017).

42. E. A. Collisson et al., Subtypes of pancreatic ductal adenocarcinoma and their differing responses to therapy. Nat Med 17, 500–503 (2011).

43. D. R. Welch, D. R. Hurst, Defining the Hallmarks of Metastasis. Cancer research 79, 3011–3027 (2019).

44. N. Muthalagu et al., Repression of the Type I Interferon pathway underlies MYC & KRAS-dependent evasion of NK & B cells in Pancreatic Ductal Adenocarcinoma. Cancer Discov, (2020).

45. R. M. Kortlever et al., Myc Cooperates with Ras by Programming Inflammation and Immune Suppression. Cell 171, 1301–1315 e1314 (2017).

46. N. M. Sodir et al., MYC Instructs and Maintains Pancreatic Adenocarcinoma Phenotype. Cancer Discov 10, 588–607 (2020).

47. K. A. Jablonski et al., Novel Markers to Delineate Murine M1 and M2 Macrophages. PLoS One 10, e0145342 (2015).

48. B. Costa-Silva et al., Pancreatic cancer exosomes initiate pre-metastatic niche formation in the liver. Nat Cell Biol 17, 816–826 (2015).

49. Y. Jia et al., IL24 and its Receptors Regulate Growth and Migration of Pancreatic Cancer Cells and Are Potential Biomarkers for IL24 Molecular Therapy. Anticancer Res 36, 1153–1163 (2016).

50. T. Kodama et al., CCL3-CCR5 axis contributes to progression of esophageal squamous cell carcinoma by promoting cell migration and invasion via Akt and ERK pathways. Lab Invest 100, 1140–1157 (2020).

51. L. Miao et al., Targeting the STING pathway in tumor-associated macrophages regulates innate immune sensing of gastric cancer cells. Theranostics 10, 498–515 (2020).

52. C. W. Steele et al., CXCR2 Inhibition Profoundly Suppresses Metastases and Augments Immunotherapy in Pancreatic Ductal Adenocarcinoma. Cancer cell 29, 832–845 (2016).

53. Y. Zhu et al., CSF1/CSF1R blockade reprograms tumor-infiltrating macrophages and improves response to T-cell checkpoint immunotherapy in pancreatic cancer models. Cancer research 74, 5057–5069 (2014).

54. Y. Zhu et al., Tissue-Resident Macrophages in Pancreatic Ductal Adenocarcinoma Originate from Embryonic Hematopoiesis and Promote Tumor Progression. Immunity 47, 323–338 e326 (2017).

55. L. Cassetta et al., Human Tumor-Associated Macrophage and Monocyte Transcriptional Landscapes Reveal Cancer-Specific Reprogramming, Biomarkers, and Therapeutic Targets. Cancer cell 35, 588–602 e510 (2019).

56. L. Cassetta, J. W. Pollard, Tumor-associated macrophages. Curr Biol 30, R246–R248 (2020).

57. P. S. Ginter et al., Tumor Microenvironment of Metastasis (TMEM) Doorways Are Restricted to the Blood Vessel Endothelium in Both Primary Breast Cancers and Their Lymph Node Metastases. Cancers (Basel) 11, (2019).

58. M. Roh-Johnson et al., Macrophage contact induces RhoA GTPase signaling to trigger tumor cell intravasation. Oncogene 33, 4203–4212 (2014).

59. C. C. Lee et al., Macrophage-secreted interleukin-35 regulates cancer cell plasticity to facilitate metastatic colonization. Nat Commun 9, 3763 (2018).

60. Q. Chen, X. H. Zhang, J. Massague, Macrophage binding to receptor VCAM-1 transmits survival signals in breast cancer cells that invade the lungs. Cancer cell 20, 538–549 (2011).

61. K. L. Aung et al., Genomics-Driven Precision Medicine for Advanced Pancreatic Cancer: Early Results from the COMPASS Trial. Clin Cancer Res 24, 1344–1354 (2018).

62. S. Turajlic, A. Sottoriva, T. Graham, C. Swanton, Resolving genetic heterogeneity in cancer. Nat Rev Genet 20, 404–416 (2019).

63. A. W. Lambert, D. R. Pattabiraman, R. A. Weinberg, Emerging Biological Principles of Metastasis. Cell 168, 670–691 (2017).

64. E. Hessmann, G. Schneider, V. Ellenrieder, J. T. Siveke, MYC in pancreatic cancer: novel mechanistic insights and their translation into therapeutic strategies. Oncogene 35, 1609–1618 (2016).

65. A. S. Farrell et al., MYC regulates ductal-neuroendocrine lineage plasticity in pancreatic ductal adenocarcinoma associated with poor outcome and chemoresistance. Nat Commun 8, 1728 (2017).

66. A. P. Makohon-Moore et al., Limited heterogeneity of known driver gene mutations among the metastases of individual patients with pancreatic cancer. Nature genetics 49, 358–366 (2017).

67. C. V. Dang, MYC on the path to cancer. Cell 149, 22–35 (2012).

68. A. V. Vaseva et al., KRAS Suppression-Induced Degradation of MYC Is Antagonized by a MEK5-ERK5 Compensatory Mechanism. Cancer cell 34, 807–822 e807 (2018).

69. J. M. Arriaga et al., A MYC and RAS co-activation signature in localized prostate cancer drives bone metastasis and castration resistance. Nature Cancer 1, 1082–1096 (2020).

70. A. Hayashi et al., Genetic and clinical correlates of entosis in pancreatic ductal adenocarcinoma. Mod Pathol, (2020).

71. T. Baslan et al., Novel insights into breast cancer copy number genetic heterogeneity revealed by single-cell genome sequencing. Elife 9, (2020).

72. B. C. Lewis, D. S. Klimstra, H. E. Varmus, The c-myc and PyMT oncogenes induce different tumor types in a somatic mouse model for pancreatic cancer. Genes Dev 17, 3127–3138 (2003).

73. F. X. Schaub et al., Pan-cancer Alterations of the MYC Oncogene and Its Proximal Network across the Cancer Genome Atlas. Cell Syst 6, 282–300 e282 (2018).

